# Piperine activates the thick filament of resting rat skeletal muscle, enhancing dynamic contractility in a fibre-type-dependent manner

**DOI:** 10.1101/2025.11.24.689918

**Authors:** Daniel Z. Kruse, Jon Herskind, Michel N. Kuehn, Annika J. Klotz, Anthony L. Hessel, Kristian Overgaard

## Abstract

The myosin-containing thick filament has recently been shown to alter its resting activation level in response to multiple diseases and therapeutics. Changes in thick filament resting activation level are caused by myosin heads transitioning between OFF and ON conformational states. Functionally, this modulation of thick filament activation level is a key regulatory step in muscle contraction and a promising therapeutic target. The availability of resting ON-state myosin heads governs dynamic contractility, which is critical to physical function and well-being. At present, there is a lack of compounds favouring this ON-state in resting skeletal muscle. Piperine is a molecule known to bind to myosin and increase submaximal isometric contractility in fast and slow skeletal muscle. Yet, effects on dynamic contractility and the underlying mechanism responsible for the observed effects in skeletal muscles remain unclear. Here, we used fibre small-angle X-ray diffraction and intact-muscle ex vivo contractility experiments to determine the effects of piperine on resting myosin structure and dynamic contractility in fast and slow rat muscles. X-ray diffraction data suggest that piperine promotes an OFF-to-ON transition of myosin in resting skeletal muscle, increasing the availability of myosin heads for force generation. Functionally, piperine substantially enhanced dynamic contractility in both muscle types, with greater improvements in slow muscle during maximal activation. These findings establish piperine as a tool to probe thick-filament activation in skeletal muscle, highlighting fibre-type–specific effects of thick-filament activation on the recruitment of the contractile reserve capacity.

**Key Points:**

– Piperine is a compound known to bind to skeletal muscle myosin and enhance isometric contractility in fast and slow muscles, but the underlying molecular mechanisms and effects on dynamic contractile function remain unknown.
– We show that piperine increases the activation level of the myosin-containing thick filament in resting fast and slow skeletal muscle, which may explain the effect of piperine on contractile function.
– Piperine substantially increases the maximal contractile power of both fast and slow skeletal muscles at low-frequency activation; however, it only enhances the maximal power in slow skeletal muscle at high-frequency activation.
– Our data reveal potentiation of dynamic contractility with fibre-type-dependent magnitudes in response to piperine-induced activation of the resting thick filament, a phenomenon that requires further investigation and may ultimately be exploited in the treatment of diseases characterised by muscle weakness.

Graphical abstract
Abstract figure legend: We investigated the effects of piperine on 1) the activation level of the resting thick filament and 2) dynamic contractility in fibres and intact slow (soleus) and fast (extensor digitorum longus, EDL) rat muscles, respectively. The activation level of the resting thick filament was assessed pre– and post-piperine incubation using small-angle X-ray diffraction. Dynamic contractility was assessed at submaximal and maximal activation levels by constructing low– and high-frequency force-velocity curves and corresponding power curves using an ex vivo contraction setup. The setup allows for simultaneous experiments on the effects of piperine and vehicle treatment in contralateral muscles. We found that piperine increased the activation level of the resting thick filament by favouring the ON-myosin state in fibres from both muscle types. In whole muscle preparations, piperine also induced substantial increases in dynamic contractility, especially in slow soleus muscle.

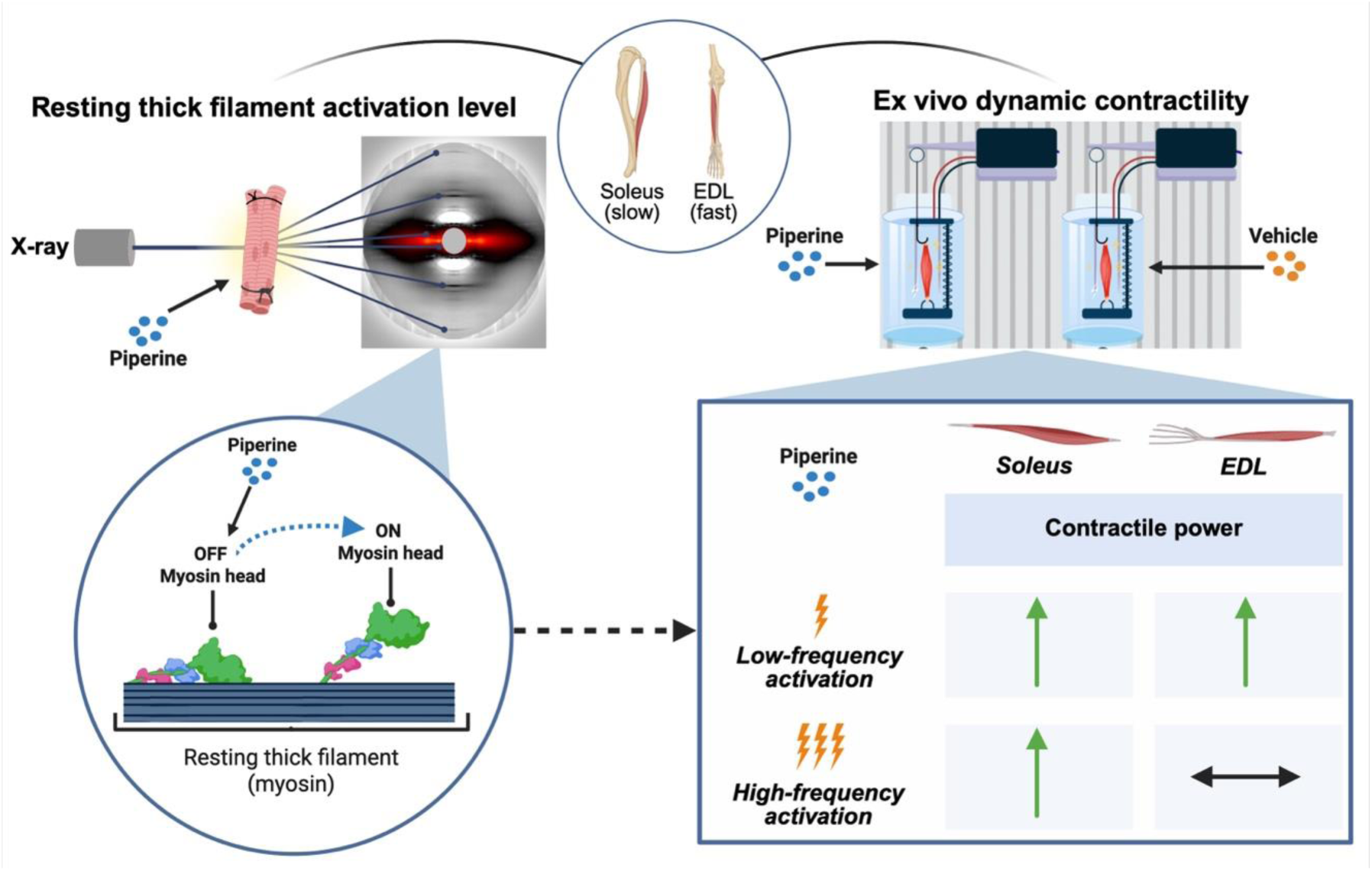

## Introduction

Recent advances in muscle physiology have highlighted the myosin-containing thick filament as an important regulator of muscle contractility (Irving, 2017; Brunello & Fusi, 2024; Hill *et al*., 2025). Therefore, the thick filament also serves as a promising therapeutic target for restoring contractile impairments in cardiac and skeletal muscle (Lehman *et al*., 2022; Claassen *et al*., 2023). In this perspective, modulation of dynamic muscle function is of particular interest, as it is paramount to physical functioning and well-being (Freitas *et al*., 2024). Resting myosin heads exist in an equilibrium between the so-called ON and OFF structural states. ON myosin heads are located near actin and are available for interaction, whereas OFF heads are tightly ordered near or within the helical tacks of the thick filament backbone (Ma & Irving, 2022). These structural states of myosin in relaxed muscle have previously been shown to regulate dynamic muscle function during contraction via modulating the propensity of each myosin to form crossbridges and thus impact performance (Piazzesi *et al*., 2007). While pharmacological small-molecules are proving to be promising thick-filament modulators in treating contractile impairments in cardiomyopathies (Olivotto *et al*., 2020; Voors *et al*., 2020; Lehman *et al*., 2022), there is a lack of molecules specifically designed to target skeletal muscle.

A potential tool for investigating thick-filament activation in skeletal muscle is the compound piperine, a natural alkaloid found in black pepper (Nogara *et al*., 2016; Lewis *et al*., 2025). Piperine has been shown to bind to the myosin regulatory light chain (RLC) (Nogara *et al*., 2016; Tolkatchev *et al*., 2018), a domain known to regulate muscle contractility (Vandenboom, 2016). Using intact rat muscles, Herskind et al. 2024 (Herskind *et al*., 2024) demonstrated that piperine increases isometric twitch force and force of unfused isometric contractions in both fast– and slow-twitch skeletal muscles. Further, piperine increased myofibrillar Ca²⁺ sensitivity, a fundamental property of muscle contraction, in both fibre types without altering resting or contraction-induced sarcoplasmic Ca²⁺ release (Herskind *et al*., 2024). Interestingly, piperine also enhances Ca²⁺ sensitivity in cardiomyocytes (Jani & Ma, 2025), and this effect was ascribed to an increase in ON myosin (Jani *et al*., 2024). Together, these studies suggest that piperine elicits a myosin-mediated contractile effect, which is shared by both cardiac and skeletal muscle; however, mechanistic explanations remain speculative.

Dynamic muscle performance in intact muscle can be described by the force–velocity curve (Alcazar *et al*., 2019), and parameters of the curve can be exploited to study contraction mechanisms. An important molecular modulator of the force–velocity curve is the availability and proportion of ON myosin heads that are ready to engage in crossbridge formation at different external loads (Piazzesi *et al*., 2007; Linari *et al*., 2015; Fusi *et al*., 2017; Brunello & Fusi, 2024; Hill *et al*., 2025). During contraction, an OFF-to-ON transition of myosin heads naturally occurs, with the speed and degree of this transition being fibre-type dependent (Gong *et al*., 2022; Hill *et al*., 2022; Zhao *et al*., 2024; Hill *et al*., 2025). This may suggest the existence of a pool of constitutively OFF myosin motors, representing a contractile reserve capacity, which seems to differ between fibre types during normal contraction. Expanding on this, it is unknown if an increased proportion of ON myosin heads in resting skeletal muscle leads to fibre-type-dependent changes in dynamic contractility. Answering these unknowns is essential to address the therapeutic potential of thick-filament activation in the treatment of debilitating myopathies (Ochala *et al*., 2010; Lindqvist *et al*., 2019; Karimi *et al*., 2024). Accordingly, in the present study, we aimed to test the hypothesis that the underlying cause of piperine’s force-potentiating effects in both fast and slow skeletal muscle is activation of the resting thick filament. Furthermore, to investigate the extent to which piperine could enhance contractile power in dynamic contractions in muscles containing predominantly fast– and slow-twitch fibres, respectively.

## Materials and Methods

### Ethical approval

No experiments were performed on live animals. Animal housing, handling, and euthanasia by cervical dislocation were performed by trained personnel according to local and European animal welfare regulations. All experiments were performed using isolated hindlimb muscles from 4-week-old female Wistar rats (Janvier Labs, LeGenest-Saint-Isle, France).

### Muscle preparations

Extensor digitorum longus (EDL) and soleus muscles were used for all experiments due to their predominance of fast– and slow-twitch fibres, respectively (Cornachione *et al*., 2011). For the determination of resting thick-filament structure by small-angle X-ray diffraction (MyoXRD) experiments, glycerol-permeabilised bundles of fibres from both EDL and soleus muscles were used (Hessel *et al*., 2024b). Intact muscles were dissected and tied to small sticks to maintain fixed length while being submerged in relaxing solution (concentrations in mM: Potassium propionate 91.2, MgCl_2_ 3.6, Na_2_-ATP 2.5, EGTA 7.0, Imidazole 20.0, and Halt^TM^ protease inhibitor cocktail 1% [Thermo Fisher Scientific, Waltham, MA, USA], pH 7.0 at room temperature). Permeabilisation was achieved by placing the muscles in relaxing solution at 4°C for 8 hours while on a rotator and then transferring them to a storage solution composed of the relaxing solution and glycerol (40/60% volume), where they were stored overnight on a rotator at 4°C. Hereafter, muscles were stored for 2-4 weeks at –20°C, before being transferred to the experimental facility on ice. For contractile experiments, intact EDL and soleus muscles were dissected while being moistened in Krebs-Ringer (KR) buffer (concentrations in mM: NaCl 122.13, KCl 2.84, CaCl_2_ 1.27, MgSO_4_ 1.19, KH_2_PO_4_ 1.19, and NaHCO_3_ 25.06; pH 7.2-7.4 at room temperature) and mounted in the experimental setup immediately after.

### Assessment of resting thick-filament structure using small-angle X-ray diffraction

Resting thick-filament structure was determined in permeabilised bundles of EDL (n = 28 bundles, 7 animals) and soleus (n = 20 bundles, 6 animals) fibres at 27°C before and after 20 min incubation in 50 µM piperine (Sigma-Aldrich, St Louis, MO, USA; cat.no. P49007) (dissolved in dimethyl sulfoxide, DMSO) (Herskind *et al*., 2024; Jani *et al*., 2024). On the experimental day, muscles were vigorously washed in cold relaxing solution to remove glycerol. Then, 2-6 mm long bundles of ∼5-15 muscle fibres were dissected, and silk suture knots (sizing 5-0) were tied at both ends. Finally, bundles were mounted onto a specially designed muscle mechanics rig for MyoXRD (Hessel *et al*., 2022) at a sarcomere length (SL) of 2.7 µm, corresponding to the optimal length of force production (L_o_) (Gong *et al*., 2022). MyoXRD patterns were collected using the small-angle instrument on the P03 / MiNaXS beamline 18ID (Petra III, DESY, Hamburg). Bundles were manually aligned and targeted with the X-ray beam using an inline camera, also allowing for visual control of the quality of samples during experiments. For each condition, the beam scanned the sample along its long length to produce ∼25-50 images with 100 ms exposure per image at a speed where no part of the sample was exposed to more than 300 ms of radiation in total, which in pilot studies produced negligible radiation damage. Line scans were separated by 50 µm displacements along the axis perpendicular to the long axis of the sample. The X-ray beam (∼0.103 nm wavelength) was focused to ∼20 μm x 20 μm on the sample, with an incident flux of 5×10^11^ photons per second. Diffraction patterns were collected by a Pilatus (2M, Detris) detector (172 μm × 172 μm pixel size). The sample-to-detector distance was 4.6-9.8 m, and real space was calculated using the 100-diffraction ring of silver behenate (d_001_ = 5.8380 nm) as a calibration standard. A subgroup of EDL bundles (n = 9 bundles) was tested at SL 2.4 and 2.7 µm before and after piperine incubation, using a passive ramp-hold protocol as previously described (data are presented in the Appendix) (Hessel *et al*., 2024a).

### Experimental setup for contractile experiments

For contractile experiments, contralateral whole muscles were placed in oxygenated chambers filled with 100 ml of standard KR buffer at 30°C and constantly perfused with 95% O_2_ and 5% CO_2._ muscles were mounted in a contractile setup (Model 305 Dual Muscle Lever, Aurora Scientific, Aurora, ON, Canada) as illustrated in Figure 1A. This setup allowed for simultaneous experiments on contralateral muscles, in which contraction force and muscle length were measured during electrically induced contractions. After mounting in chambers, muscles were stimulated to elicit a brief isometric tetanus to test the integrity of the knots, and then they rested for 30 min. Muscles were stretched to the length of optimal isometric force production (L_o_) determined using triplet stimulation, and thereafter the muscles rested for 20 min before the experiments began. Contractile forces were checked during the initial protocol to assess the stability of the muscle preparations. Isometric and dynamic muscle contractions were evoked by electrical stimulation with a computer-controlled lever arm imposing various external loads (DMC 5.500, Aurora Scientific). All contractions were induced by supramaximal electrical stimulation via field stimulation by two platinum plate electrodes (pulse duration = 0.2 s; 1 A; 16 V/cm^2^) (Jakobsgaard *et al*., 2022).

**Figure 1.**
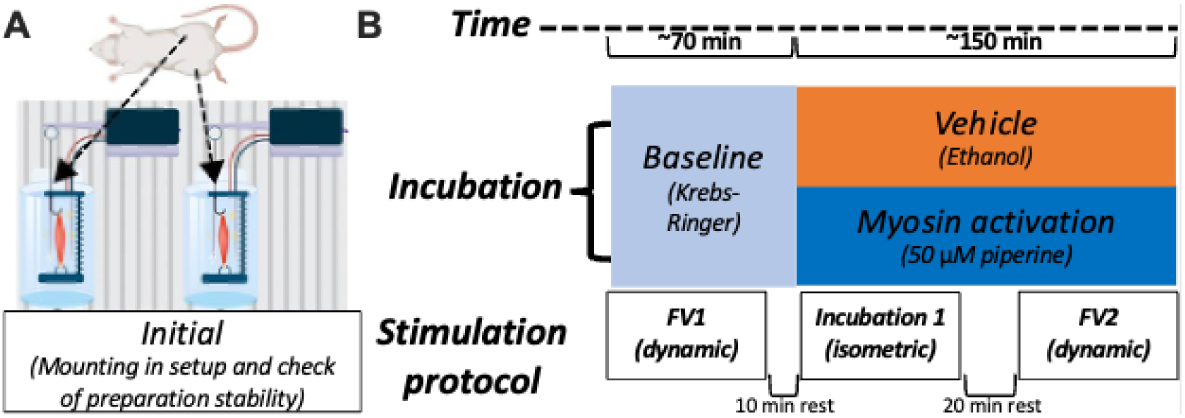
Experimental setup and main protocol for whole-muscle contractility experiments. A, contralateral pairs of intact fast (extensor digitorum longus, EDL) or slow-twitch (soleus) rat skeletal muscles were dissected and mounted in the illustrated setup. B, the main experimental protocol used to test the isolated effects of piperine (versus vehicle) on dynamic contractility. Dynamic contractility was assessed by performing force-velocity (FV) tests at baseline (FV1) and after a 1-hour incubation (FV2). The illustrations are made using Biorender.com and Microsoft PowerPoint.

### Main experimental protocol for contractile experiments

The main experimental protocol (Fig. 1B) compared the isolated effects of vehicle and 50 µM piperine treatment on dynamic contractility in contralateral pairs of fast (EDL, n = 6 animals, bodyweight 111 ± 8 g) and slow (soleus, n = 6 animals, bodyweight 91 ± 16 g) rat muscles. Dynamic muscle function was assessed by performing force-velocity tests at baseline (FV1) and after a 60 min incubation (FV2). Isometric contractility was monitored during the 60 min incubation by twitch (2 Hz), unfused tetanic (EDL, 50 Hz for 0.3 s; soleus, 20 Hz for 2 s), and fused tetanic (EDL, 150 Hz for 0.2 s; soleus, 80 Hz for 2 s) contractions induced every 20^th^ and 10^th^ min for EDL and soleus muscles, respectively. These three isometric contraction types were tested due to their distinct underlying features of intracellular Ca^2+^ release, summation of forces from subsequent action potentials, and thick-filament activation level (Brunello & Fusi, 2024; Hill *et al*., 2025). Piperine was dissolved in ethanol 96%, with ethanol (vehicle) making up 0.1% of the total buffer volume.

### Assessment of low– and high-frequency dynamic muscle function during force-velocity testing

Isotonic low– and high-frequency (EDL, 60 Hz and 150 Hz; soleus, 20 Hz and 80 Hz, respectively) force-velocity curves were determined using the afterload method, as previously described (Kristensen *et al*., 2019; Kristensen *et al*., 2020). Each force-velocity curve was determined by measuring the maximal shortening velocity (mm/s) against seven external load levels (in this order: 15, 80, 27.5, 40, 5, 100, and 60% of maximal low– and high-frequency isometric force). To allow for satisfying recording of length changes and shortening velocities during force-velocity (FV) testing, contractions were performed using varying durations of stimulation trains at the various percentages of isometric maximum force. Specifically, stimulation trains for EDL were 0.3 s for contractions at 100% (isometric) and 0.2 s for all other contractions. For the soleus, stimulation trains were 0.6 s for low– and high-frequency contractions from 5-27.5% and 5-80%, respectively, 1.2 s for low-frequency contractions from 40-80%, and 2 s for low– and high-frequency at 100% (see also exemplary force– and length traces in Figure A2). Contractions at subsequent loads were interspersed by 10 min of rest, and low– and high-frequency contractions at the same relative loads were interspersed by 5 s of rest.

### Assessment of the overall rate of crossbridge cycling

As a separate experiment, we determined the effects of 50 µM piperine (versus vehicle) on the rate of actomyosin crossbridge cycling at L_o_ in contralateral fast (EDL, n = 7 animals, bodyweight 107 ± 15 g) and slow (soleus, n = 4 animals, bodyweight 108 ± 6 g) rat muscles. The rate of force redevelopment (K_tr_) was used to assess the overall crossbridge cycling rate under maximal Ca²⁺ conditions. Experiments were performed using the same setup and incubations (60 min) as described in Figure 1. K_tr_ was experimentally determined in both muscle types, as previously described (Kristensen *et al*., 2019), every 15^th^ minute during a 1-hour incubation. Briefly, fused tetani were induced (EDL, 150 Hz for 0.46 s; soleus, 80 Hz for 1.6 s). As the tetanic isometric force was achieved, muscles were allowed to perform unloaded shortening for 10 ms, while maintaining tetanic stimulation. Thereafter, muscles were rapidly re-stretched back to L_o_ (2 ms) and allowed to redevelop tetanic isometric force. K_tr_ was estimated by fitting the force data from the redevelopment of isometric force (mean r^2^ ± SD for curve-fit: EDL 0.94 ± 0.05; soleus 0.97 ± 0.01) by the following equation:

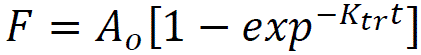

with A_o_ being the force at the plateau of the force redevelopment

### Analysis of images from MyoXRD experiments

Initially, MyoXRD patterns were reduced and prepared for analysis in a custom-made program (Bulb, Accelerated Muscle Biotechnologies, Boston, USA). After, images were analysed using the MuscleX open-source data reduction and analysis package (BioCAT, Argonne National Laboratory). In brief, images in a line scan were aligned, median-merged together, and quadrant-folded. The intensities of reflections were normalised across samples by the X-ray intensity collected at the beamline beamstop. The Equator routine of MuscleX was used to calculate the equatorial intensity ratio (I_1.1_/I_1.0_) and the inter-thick-filament lattice spacing (d_10_). To determine the equatorial intensities (I_1.0_ and I_1.1_), the intensities along the equatorial axis of the image were integrated (± 0.005 nm^−1^ on either side of the equatorial axis). This was followed by 1.0 and 1.1 fitting Gaussian curves to the 1.0 and 1.1 reflections, respectively. Spacings and intensities were calculated via the maximum Gaussian peak and area under the Gaussian curve, respectively. Spacings of the meridional M3 and M6 reflections (S_M3_ and S_M6_) were extracted by the projection-traces routine in MuscleX. The intensity of the M3 reflection (I_M3_) was calculated by integrating in the reciprocal radial range ∼0 ≤ R ≤ 0.032 nm^−1^ for the M3 reflections. Hereafter, integration limits on the median axis were 0.066 ≤ R ≤ 0.074 nm^−1^ and 0.136 ≤ R ≤ 0.144 nm^−1^ for the M3 and M6, respectively. The spacing and intensity of each reflection were calculated automatically from these. Gaussian fits with fit errors above 10% were excluded.

### Analysis of data from contractile experiments

All contractile data (force and length) were recorded at 2000 Hz and filtered using a Butterworth filter with a cutoff frequency of 30 Hz, except for the K_tr_ data, which had a cutoff frequency of 200 Hz. Data was analysed using custom-made MATLAB software (MATLAB R2024a, MathWorks Inc.) (Kristensen *et al*., 2019). Force kinetics and shortening velocity were measured over a 0.01-s average. Isometric contractions were analysed for the following outcomes: peak active force (peak total force – passive tension) (mN), peak rate of force development (mN/ms), and relaxation time (delta time in ms from 90% to 10% of the peak force). During dynamic contractions, the contractile force and the simultaneous shortening velocity were analysed. The contractile force (mN) during shortening was analysed as the average active force at the force plateau during shortening. Maximal shortening velocity (mm/s) was analysed as the largest negative change in muscle length during a 0.01-s time interval. From measured force and shortening-velocity values, force-velocity curves were fitted (mean R^2^ > 0.98) using Hill’s force equation (1), and the corresponding power curves (force × velocity) were fitted using Hill’s power equation (2):

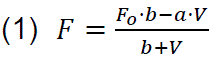

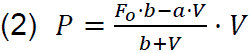

In the above equations, F_o_ is the measured maximal isometric force, V is the shortening velocity, and a and b are constants. The maximal unloaded shortening velocity (V_max_) (mm/s) and maximal isometric force production (F_max_) (mN) were estimated from the extrapolation of these curves. Likewise, the curvature of the force-velocity curve (a/F_o_) (a.u.) and the maximal power production (P_max_) (mJ/s) were estimated.

### Statistical analysis

Descriptive values are presented as mean ± standard deviation (SD), while differences (absolute and percentage) are presented as mean difference followed by a 95% confidence interval. A significance level of 0.05 was used, and all statistical analyses and figures were made using GraphPad Prism 10.2 (GraphPad Software, La Jolla, CA, USA). Data were assessed for normality by visual inspection of quantile-quantile (QQ) plots, and assumptions of equal standard deviations were tested by F-tests. To meet assumptions of normality, some MyoXRD outcomes (EDL: I_1.1_/I_1.0_, S_M3_, and I_M3_; soleus: I_1.1_/I_1.0_) were log-transformed for statistical analysis. Changes in MyoXRD outcomes were analysed using repeated measures (RM) ANOVAs comparing baseline and piperine values. Changes in isometric and dynamic contractile function were analysed by RM ANOVAs comparing relative changes from baseline between muscle pairs treated with piperine and vehicle. For K_tr_ measurements, relative changes from baseline at all time points were compared between paired piperine– and vehicle-treated muscles using a two-way RM ANOVA with Tukey’s correction.

## Results

### Piperine increases the thick filament activation level in resting slow– and fast-twitch fibres

To determine the currently unknown molecular changes that can explain the previously reported changes in isometric contractility of both slow and fast skeletal muscle (Herskind *et al*., 2024), we assessed the activation level of the resting thick filament before and after piperine treatment using MyoXRD (Fig. 2A and B).

**Figure 2.**
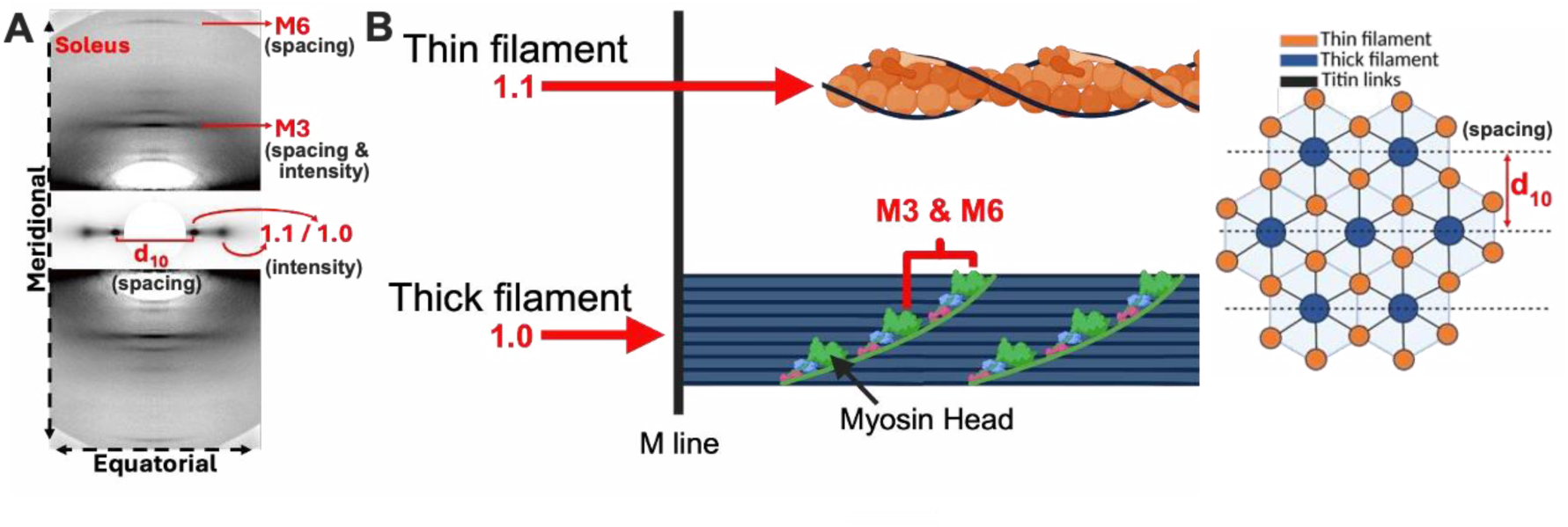
Schematic of the resting thick filament structure assessed by fibre small-angle X-ray diffraction (MyoXRD) experiments. A, a representative MyoXRD pattern from one soleus bundle with reflections originating from longitudinally (meridional) and transversely (equatorial) repeating structures. In the representative image, it is stated whether the intensity (molecular density) and/or spacing (molecular distances) of the reflection was extracted. B, an illustration of the sarcomeric structures from which the diffractions arise. Red lines show the primary origin of the reflection. The following outcomes were extracted: the equatorial intensity ratio (I_1.1_ / I_1.0_), the spacings of the M3 and M6 reflections (S_M3_ and S_M6_), the intensity of the M3 reflection (I_M3_), and the inter-thick-filament lattice spacing (d_10_) (illustrated in the top-right corner (Hessel et al., 2024a). The 1.0 and 1.1 reflections arise from the myosin-containing thick filament (blue) and the actin-containing thin filament (orange), respectively. The M3 and M6 reflections primarily arise from the repeating arrays of myosin heads (green). The illustration in B is made using Biorender.com

*Fast-twitch fibres*. — The equatorial intensity ratio (I_1.1_/I_1.0_) arises from the mass distribution between the thin (I_1.1_) and thick (I_1.0_) filaments. Piperine increased the I_1.1_/I_1.0_ in fast-twitch EDL fibres compared to baseline (0.58 ± 0.21 (mean ± SD) vs. 0.94 ± 0.55 a.u., P = 0.002) (Fig. 3A), indicating a mass movement (i.e., myosin heads) from the thick filaments towards the thin filaments. The spacing and intensity of the meridional M3 reflection (S_M3_ and I_M3_) primarily arise from the average distance and general ordering, respectively, of consecutive myosin heads along the thick filament. Hence, increased spacing and less intensity of the M3 are interpreted as an increased proportion of ON myosin heads (Ma & Irving, 2022). Surprisingly, the S_M3_ (Fig. 3B) and I_M3_ (Fig. 3C) did not change significantly (S_M3_: 14.39 ± 0.03 vs. 14.42 ± 0.04 nm, P = 0.139; I_M3_: 605 ± 166 vs. 536 ± 180 a.u., P = 0.208) in response to piperine. The spacing of the meridional M6 reflection (S_M6_) primarily arises from the distance between the repeats of folded-back myosin heads, with minor contributions from other structures within the thick filament (Hessel, 2025; Koubassova *et al*., 2025). The S_M6_ (Fig. 3D) increased slightly in response to piperine (7.23 ± 0.01 vs. 7.24 ± 0.01 nm, P = 0.016), indicative of myosin heads leaving their anchored position on the thick filament. The inter-thick-filament lattice spacing (d_10_; Fig. 3E) relates to the compression of the sarcomere and was unaffected by piperine (39.9 ± 2.3 vs. 39.2 ± 1.8 nm, P = 0.964). Finally, the intensity of the first-order myosin-layer line (I_MLL1_), another marker of thick-filament activation, arising from the helical repeat of myosin crowns (Hessel, 2025; Koubassova *et al*., 2025), indicates myosin head ON/OFF states. The I_MLL1_ decreased after piperine incubation, supporting decreased ordering of myosin heads (I_MLL1_: 1663 ± 716 vs. 999 ± 523 a.u., P = 0.181; Table A1). In summary, our data indicate that piperine induced activation of the thick filament in resting fast-twitch EDL fibres (Fig. 3F; Table A1) (Linari *et al*., 2015; Ma *et al*., 2021).

**Figure 3.**
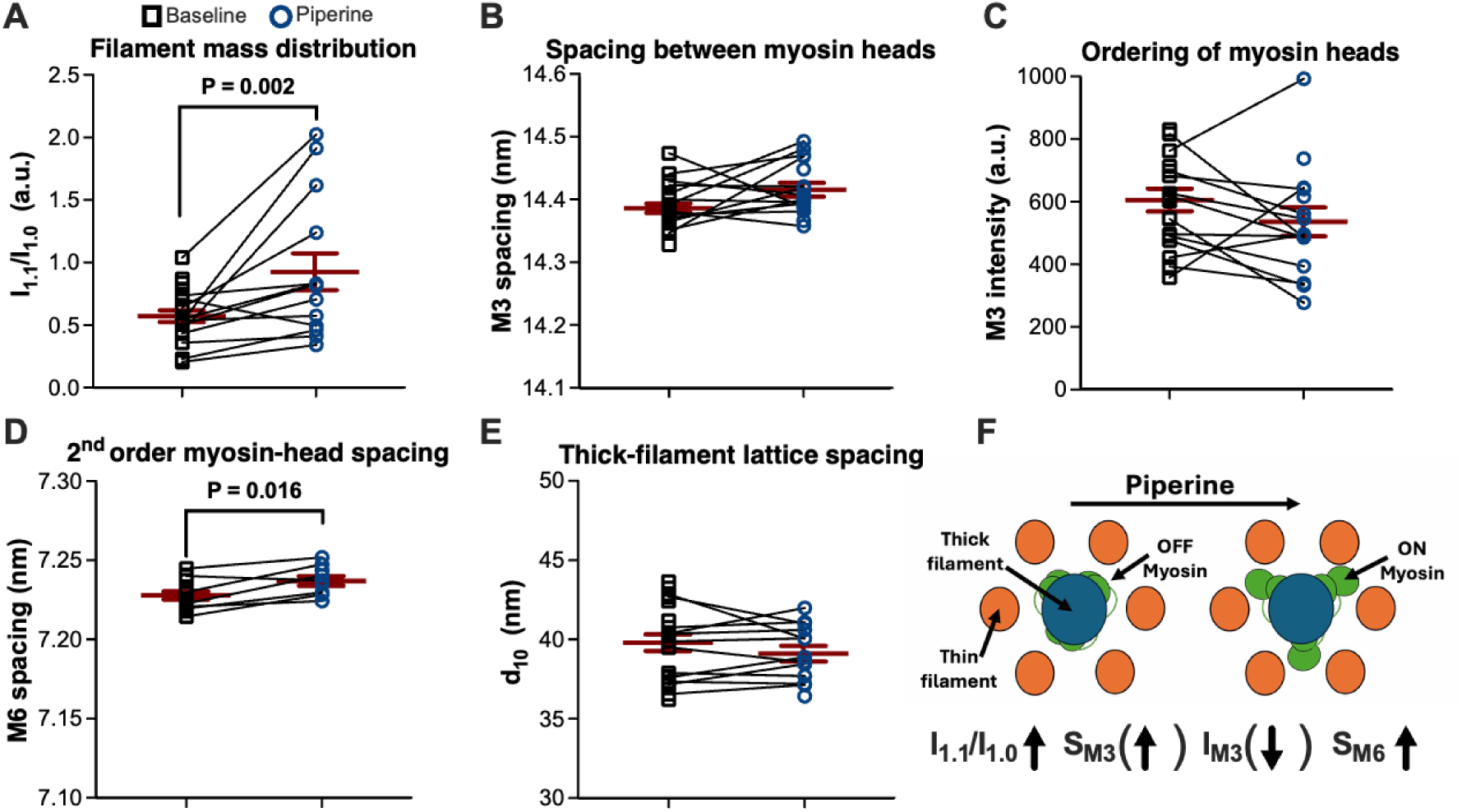
Effects of piperine on resting thick filament structure in bundles of fast (extensor digitorum longus, EDL) rat skeletal fibres. This figure shows values from individual bundles from testing at baseline (black squares) and after 20-min exposure to piperine (blue circles). Repeated measurements from the same bundle are shown by connecting lines, and group means ± SEM are shown as red lines with error bars. The following outcomes are shown: the equatorial intensity ratio (I_1.1_ / I_1.0_) (A), the spacing and intensity of the M3 reflection (S_M3_ and I_M3_) (B and C), the spacing of the M6 reflection (S_M6_) (D), and the inter-thick-filament lattice spacing (d_10_) (E). F, a scheme illustrating our interpretation of the changes induced by piperine. Vertical arrows indicate that the change was statistically significant, and vertical arrows in parentheses indicate the numerical direction of non-significant findings. P-values show comparisons of baseline and piperine values for 9-21 bundles. The illustration in F is made using Microsoft PowerPoint.

*Slow-twitch fibres*. — The slow-twitch soleus fibres showed similar but more consistent effects of piperine on resting thick-filament structure. In response to piperine, the I_1.1_/I_1.0_ showed a marked increase (0.56 ± 0.26 vs. 1.31 ± 0.85 a.u., P < 0.001; Fig. 4A), indicating a movement of mass from the thick filaments towards the thin filaments. Further, piperine increased the S_M3_ (14.40 ± 0.03 vs. 14.43 ± 0.03 nm, P = 0.009; Fig. 4B) and decreased the I_M3_ (495 ± 102 vs. 398 ± 141 a.u., P = 0.019; Fig. 4C), indicating recruitment of myosin heads from their resting position on the thick filament backbone. The S_M6_ showed a non-significant increase (7.23 ± 0.01 vs. 7.24 ± 0.01 nm, P = 0.148; Fig. 4D). Once again, the d_10_ was unaffected by piperine (39.8 ± 2.2 vs. 40.1 ± 1.9 nm, P = 0.070; Fig. 4E), suggesting no effects on sarcomere compression. Finally, the I_MLL1_ decreased markedly in response to piperine (1544 ± 665 vs. 681 ± 190 a.u., P = 0.014; Table A1). Hence, our data consistently showed a strong OFF-to-ON transition of the thick filament in the soleus fibres in response to piperine (Fig. 4F; Table A1). Taken together, our data indicate that piperine activated the thick filament in resting fibres of both types of skeletal muscle, manifesting as an OFF-to-ON signature across the diffraction outcomes.

**Figure 4.**
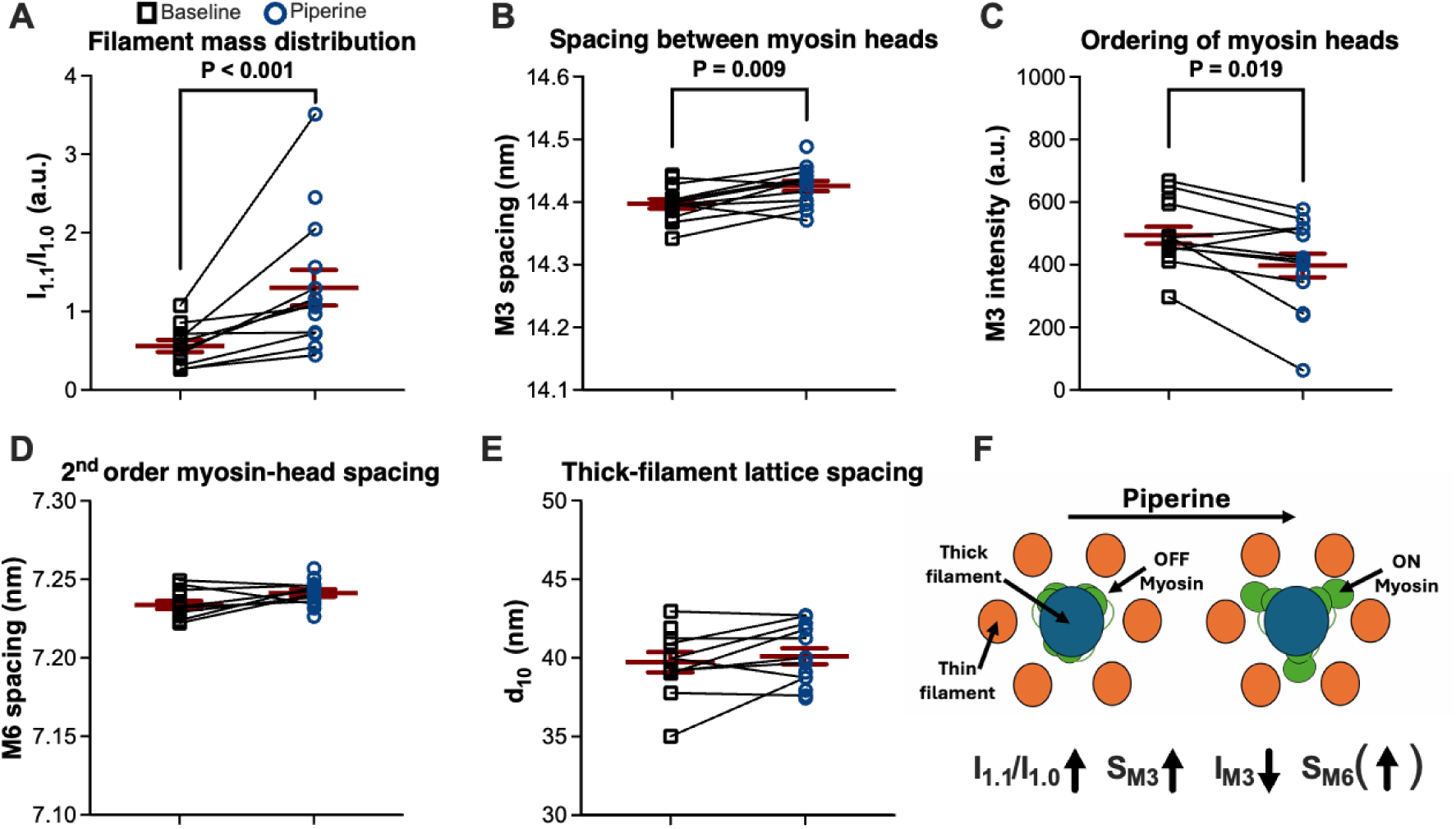
Effects of piperine on resting thick filament structure in bundles of slow (soleus) rat skeletal fibres. This figure shows values from individual bundles from testing at baseline (black squares) and after 20-min exposure to piperine (blue circles). Repeated measurements from the same bundle are shown by connecting lines, and group means ± SEM are shown as red lines with error bars. The following outcomes are shown: the equatorial intensity ratio (I_1.1_ / I_1.0_) (A), the spacing and intensity of the M3 reflection (S_M3_ and I_M3_) (B and C), the spacing of the M6 reflection (S_M6_) (D), and the inter-thick-filament lattice spacing (d_10_) (E). F, a scheme illustrating our interpretation of the changes induced by piperine. Vertical arrows indicate that the change was statistically significant, and vertical arrows in parentheses indicate the numerical direction of non-significant findings. P-values show comparisons of baseline and piperine values for 11-14 bundles. The illustration in F is made using Microsoft PowerPoint.

### Whole-muscle isometric contractile experiments demonstrate contraction-type-dependent effects of piperine

From prior literature (Herskind *et al*., 2024), we expected that isometric contractility would be enhanced by piperine, which we explored in isolated whole muscles during a 60-min incubation. We report differences between muscles treated with piperine and vehicle, expressed relative to the baseline level.

*Fast-twitch muscles*. — In EDL muscles, piperine increased twitch (Fig. 5A) and unfused tetanic (Fig. 5B) forces, while not affecting fused tetanic force (Fig. 5C). After 60 min of incubation, piperine increased twitch and unfused tetanic force by 36% [29:43] (mean [95%CI]) (P < 0.001) and 38% [31:46] (P < 0.001), respectively. At the same time, the maximal rate of force development (RFD) (Table A2) of the twitch, unfused, and fused tetanic contractions increased by 34% [28:41] (P < 0.001), 35% [27:43] (P < 0.001), and 2.0% [1.1:2.8] (P < 0.001), respectively. Finally, 60 min of piperine treatment increased the force relaxation time (RT) (Table A2) of the twitch by 3.5% [1.1:5.8] (P < 0.008). In contrast, piperine decreased the RT of unfused contractions by 6.0% [3:9.1] (P < 0.001), while the fused contractions did not show any consistent changes in RT (2.6% [-0.2:5.4], P = 0.144).

**Figure 5.**
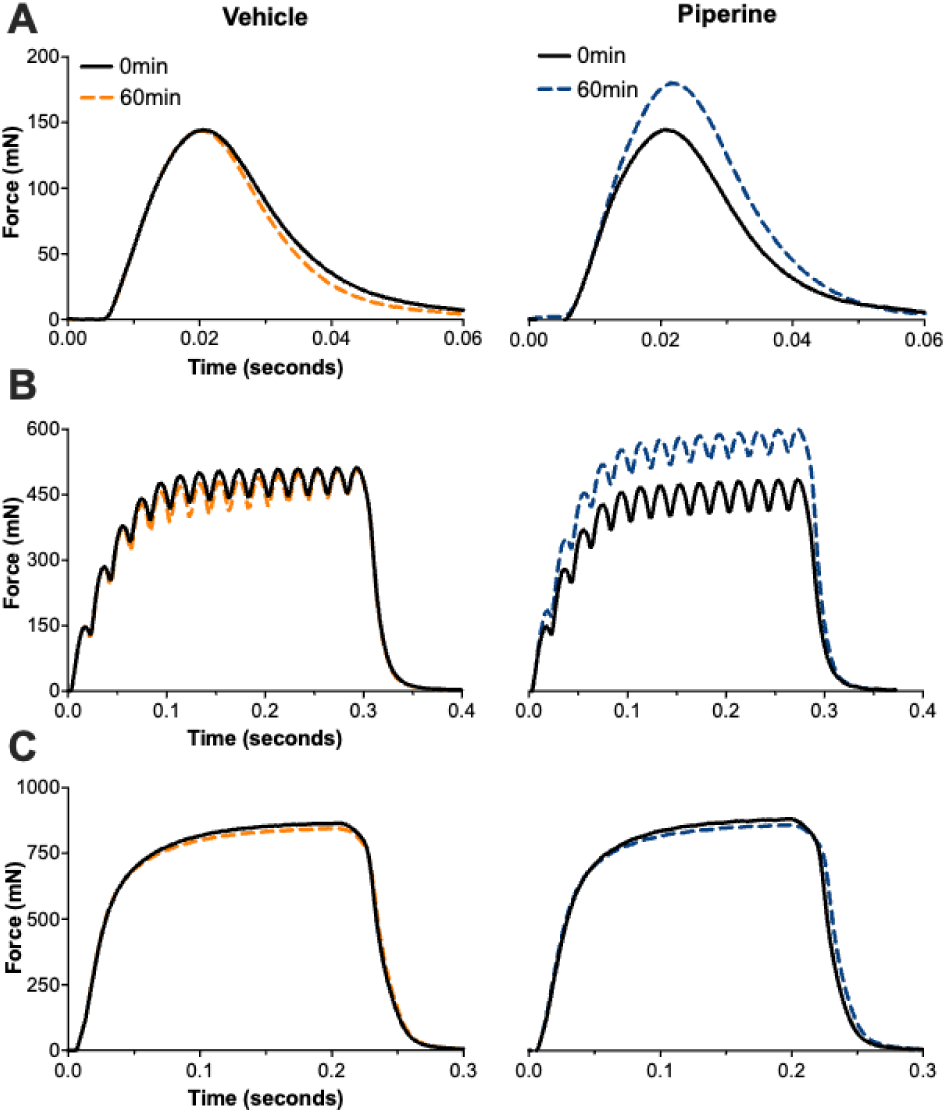
Effects of piperine on isometric contractility of intact fast (extensor digitorum longus, EDL) rat skeletal muscles. This figure shows exemplary force traces from EDL muscles exposed to vehicle and piperine at the start (0 min) and end (60 min) of incubations. Force traces (n = 1 muscle per treatment) from twitch (2 Hz) (A), unfused tetanic (50 Hz) (B), and fused tetanic (150 Hz) (C) contractions are shown.

*Slow-twitch muscles*. — Similar to the EDL muscles, the twitch (Fig. 6A) and unfused tetanic (Fig. 6B) forces of the soleus muscles increased in response to piperine, while the fused tetanic force was unaffected (Fig. 6C). In the soleus, piperine increased twitch and unfused force by 43% [37:49] (P < 0.001) and 21% [16:27] (P < 0.001), respectively. Twitch and unfused tetanic RFD of the soleus muscles (Table A3) increased by 32% [27:37] (P < 0.001) and 37% [31:42] (P < 0.001), respectively. Notably, RFD of fused tetanic contractions in soleus muscles increased substantially by 16% [13:19] (P < 0.001). Also, piperine increased soleus twitch RT (Table A3) by 5.3% [0.5:10.2] (P < 0.035), decreased unfused tetanic RT by 7.3% [5:9.7] (P < 0.001), and did not show consistent changes in RT of fused contractions. Finally, in an additional incubation phase (Fig. A1), we showed that the effects of piperine were both reversible and reproducible (EDL, Table A2; soleus, Table A3).

**Figure 6.**
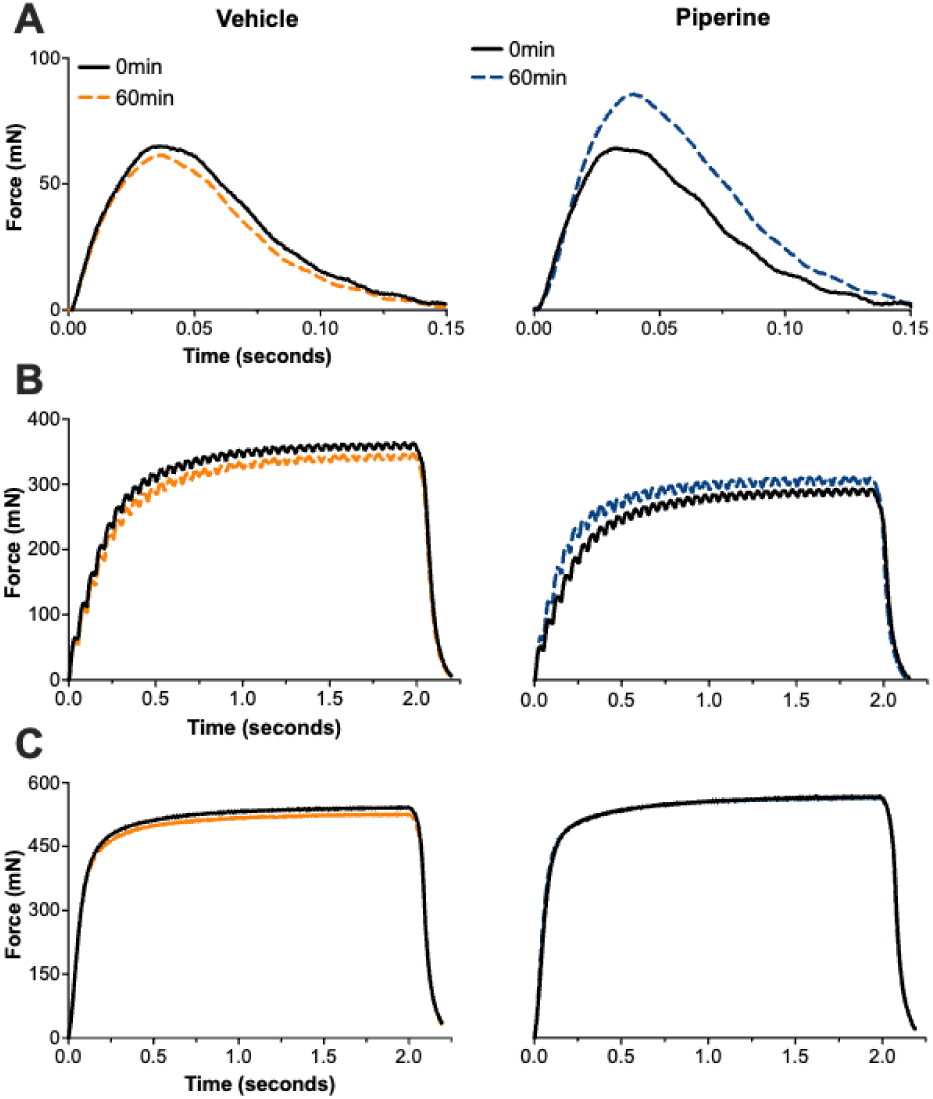
Effects of piperine on isometric muscle function of intact slow (soleus) rat skeletal muscles. This figure shows exemplary force traces from soleus muscles exposed to vehicle and piperine at the start (0 min) and end (60 min) of incubations. Force traces (n = 1 muscle per treatment) from twitch (2 Hz) (A), unfused tetanic (20 Hz) (B), and fused tetanic (80 Hz) (C) contractions are shown.

### Thick-filament activation by piperine enhances low– and high-frequency dynamic contractility, with more pronounced effects in slow-twitch muscle

Muscles working *in vivo* will rarely experience activation frequencies leading to maximal fused tetanic contraction; therefore, a comprehensive assessment of muscle function must also include testing at submaximal activation frequencies. The contractile power is largely determined by the inverse relationship between contraction force and shortening velocity; however, the curvature of the force-velocity curve also modifies power. Accordingly, force-velocity curves were determined at fibre-type-specific low– and high-frequency stimulation protocols. Below, we show exemplary force-velocity contractions (Fig. 7A) along with EDL and soleus force-velocity curves and power curves from baseline (FV1) testing (Fig. 7B and C). We report differences between muscles treated with piperine and vehicle (FV2) expressed relative to baseline (FV1). Absolute values from force-velocity testing are shown in Tables A4 and A5.

**Figure 7.**
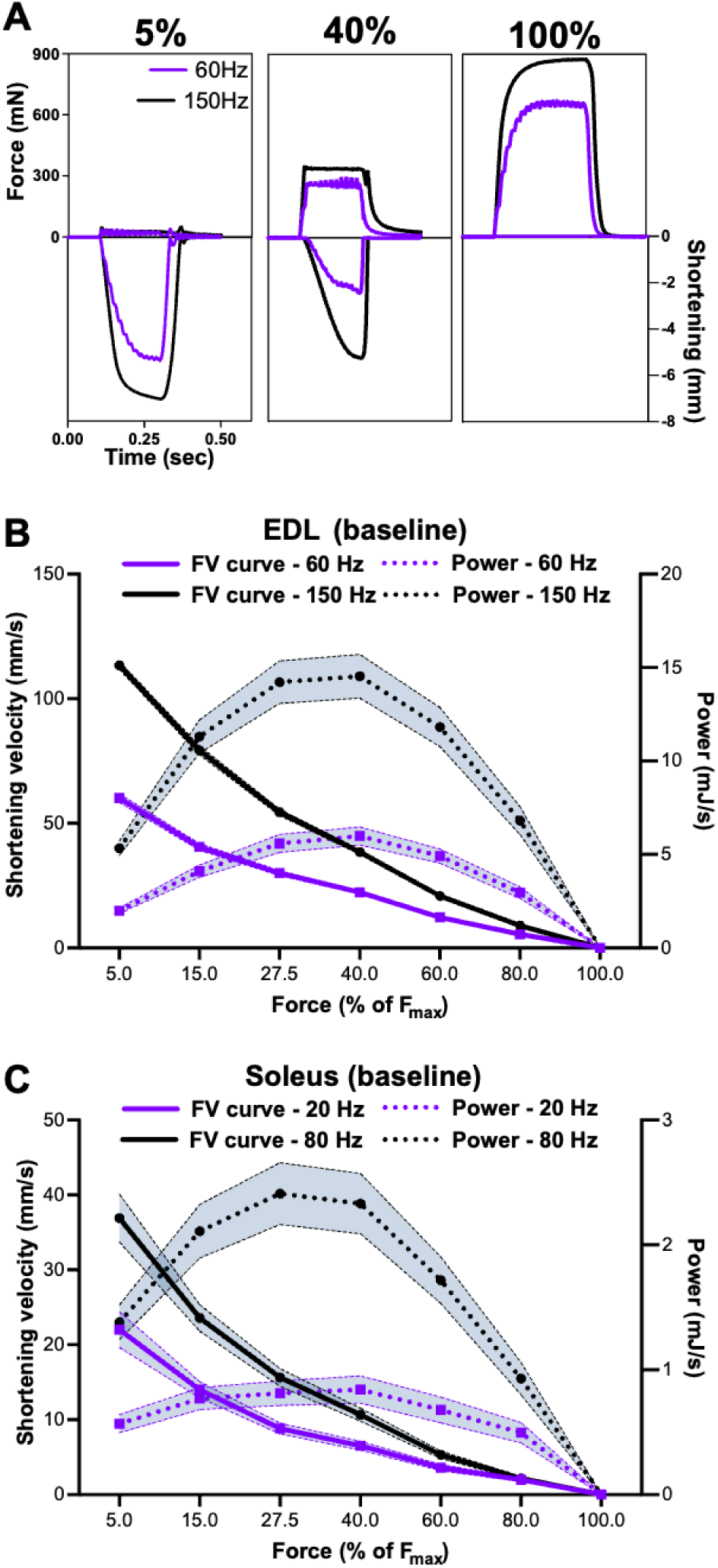
Assessment of low– and high-frequency dynamic contractility by force-velocity testing. A, exemplary low-frequency (purple) and high-frequency (black) contractions (n = 1 extensor digitorum longus (EDL) muscle) at external loads of 5, 40, and 100% of maximal low-and high-frequency isometric force (F_max_). Corresponding force and shortening velocities were used to construct force-velocity (FV) and power curves. B and C, baseline (FV1) low-frequency (purple squares) and high-frequency (black circles) EDL and soleus force-velocity (solid lines) and power (dotted lines) curves, respectively (mean with shades showing SEM). n = 6 muscles of each type.

*Low-frequency activation*. — Piperine induced marked and highly consistent increases in low-frequency dynamic contractility in both muscle types. In the EDL (Fig. 8A), piperine increased low-frequency V_max_ and P_max_ by 25% [23:28] (P < 0.001) and 36% [31:40] (P < 0.001), respectively. Also, the curvature of the force-velocity curve increased (a/F_o_ decreased) by 36% [27:46] (P < 0.001). In the soleus (Fig. 8B), piperine increased low-frequency V_max_ and P_max_ by 20% [12:28] (P = 0.005) and 54% [46:62] (P < 0.001), respectively, with no effect on curvature (P = 0.277).

**Figure 8.**
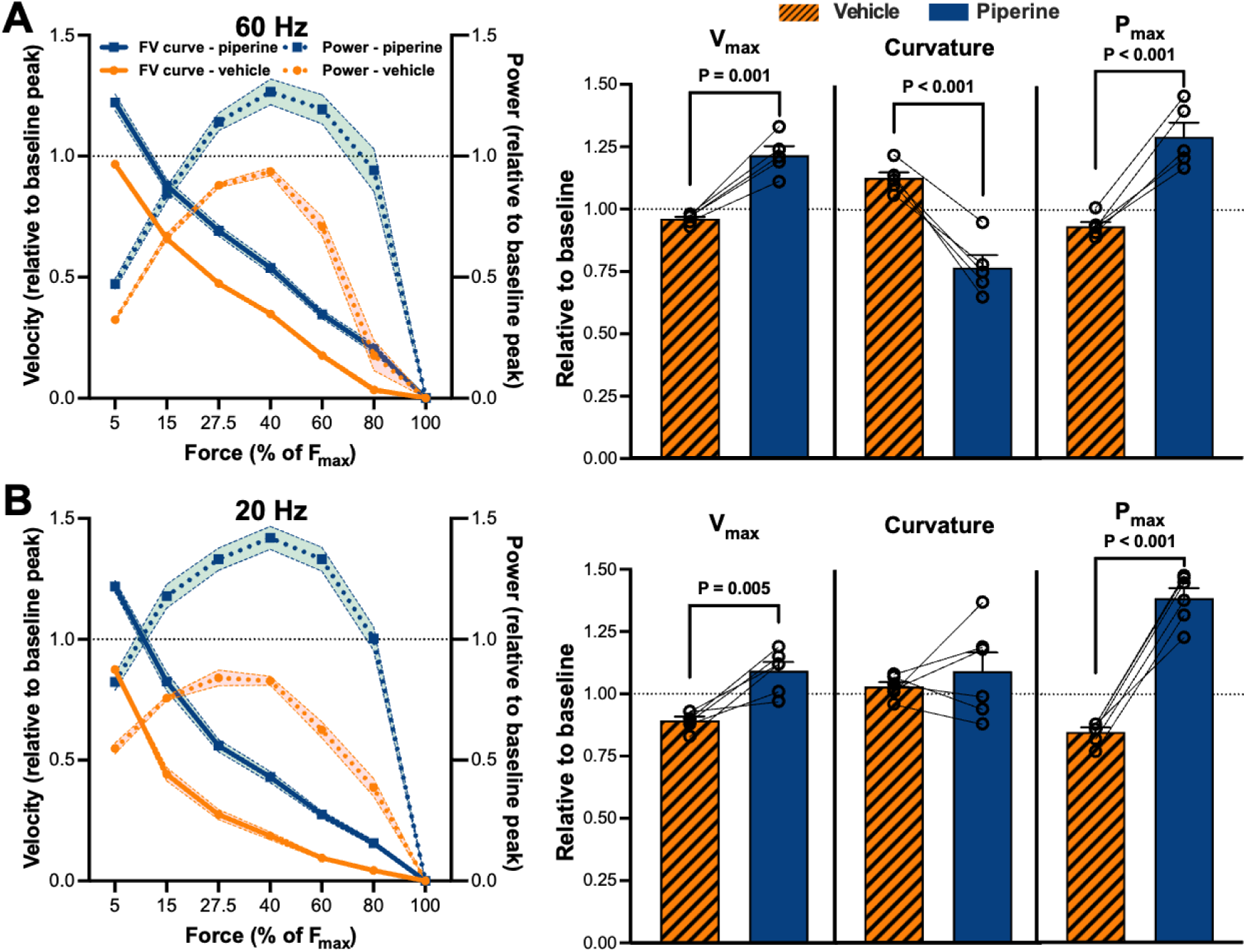
Effects of piperine on low-frequency dynamic contractility of fast (extensor digitorum longus, EDL) and slow (soleus) rat skeletal muscles. Curves on the left show low-frequency EDL (60 Hz) (A) and soleus (20 Hz) (B) force-velocity (FV) curves (solid lines) and power curves (dotted lines) after contralateral piperine (blue squares) and vehicle (orange circles) treatments. Values are normalised to baseline values and show group means, with shades indicating SEM. Bars on the right show corresponding baseline-normalised Hill-equation estimates of maximal unloaded shortening velocity (V_max_), curvature of the force-velocity curve (a/F_o_, which is inversely related to curvature), and maximal power output (P_max_) as mean ± SEM. Circles and connecting lines represent the values of individual muscles and their contralateral pairings, respectively. P-values are from comparisons of changes from baseline for piperine and vehicle. For EDL, n = 5 for piperine treatment and n = 6 for vehicle treatment. For soleus muscles, n = 6 for each treatment.

*High-frequency activation*. — Interestingly, piperine also showed effects on the high-frequency force-velocity curves in both muscle types. In the EDL (Fig. 9A), piperine increased V_max_ by 5.5% [2.6:8.3] (P = 0.001) and increased curvature (a/F_o_ decreased) by 13% [4:22] (P = 0.013), while not affecting P_max_ (P = 0.318). In contrast, the soleus muscles (Fig. 9B) exclusively showed positive effects in response to piperine, as V_max_ and P_max_ increased by respectively 9% [6:12] (P = 0.001) and 11% [7:14] (P = 0.002) without affecting curvature (P = 0.131). Finally, in an additional force-velocity test (FV3) (Fig. A1), the effects of piperine on dynamic muscle function were shown to be both reversible and reproducible (Tables A4 and A5).

**Figure 9.**
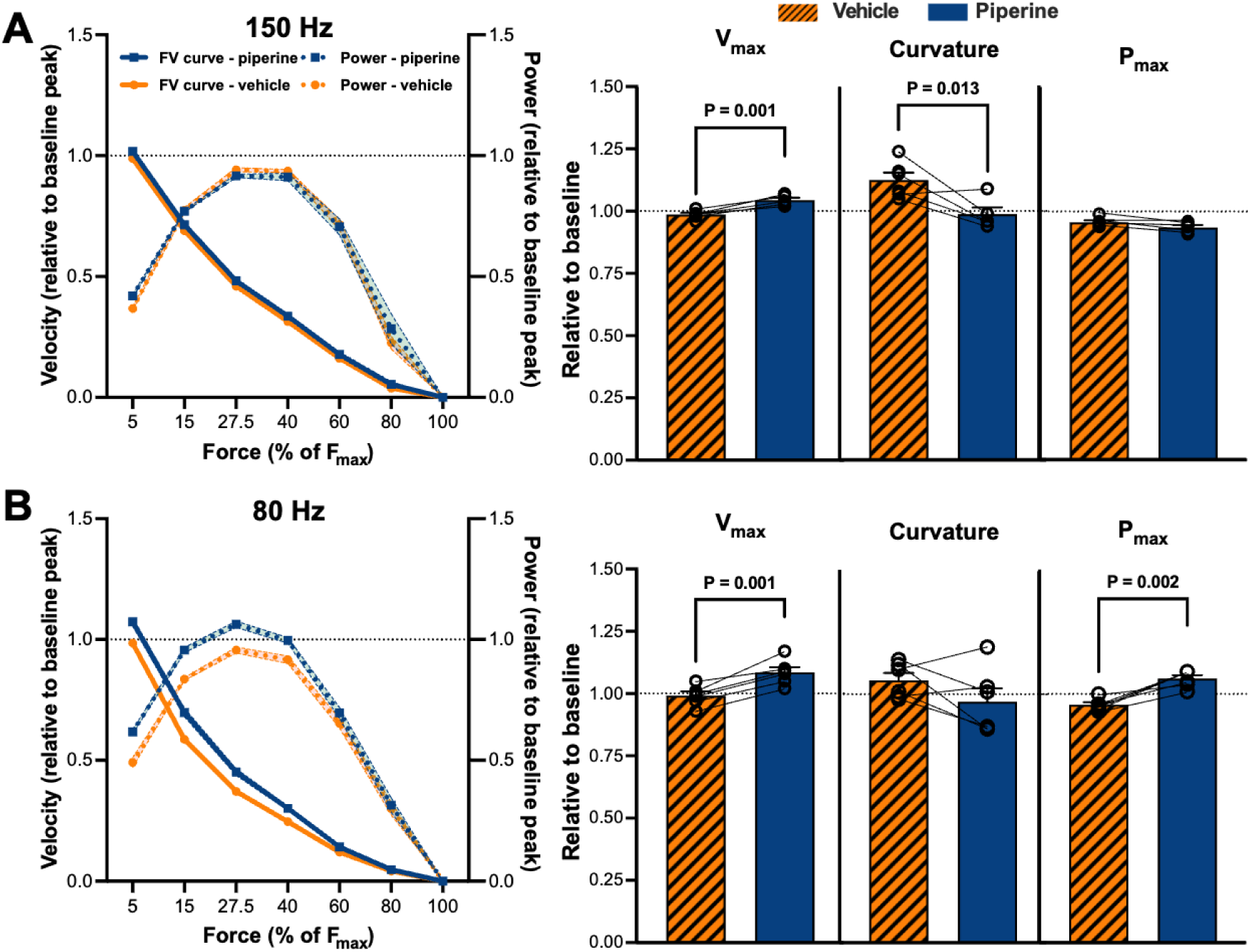
Effects of piperine on high-frequency dynamic contractility of fast (extensor digitorum longus, EDL) and slow (soleus) rat skeletal muscles. Curves on the left show high-frequency EDL (150 Hz) (A) and soleus (80 Hz) (B) force-velocity (FV) curves (solid lines) and power curves (dotted lines) after contralateral piperine (blue squares) and vehicle (orange circles) treatments. Values are normalised to baseline values and show group means, with shades indicating SEM. Bars on the right show corresponding baseline-normalised Hill-equation estimates of maximal unloaded shortening velocity (V_max_), curvature of the force-velocity curve (a/F_o_, which is inversely related to curvature), and maximal power output (P_max_) as mean ± SEM. Circles and connecting lines represent the values of individual muscles and their contralateral pairings, respectively. P-values are from comparisons of changes from baseline for piperine and vehicle. For EDL, n = 5 for piperine treatment and n = 6 for vehicle treatment. For soleus muscles, n = 6 for each treatment.

### Piperine selectively increases the overall rate of crossbridge cycling in the fast muscle type

As a proxy measure of the overall actomyosin crossbridge cycling rate, we determined K_tr_ during a 60 min incubation (Fig. 10A and B). Determination of K_tr_ allows for assessment of isometric force development under circumstances where 1) intracellular calcium levels are maximal from the beginning due to continuous electrical stimulation, and 2) the number of attached crossbridges at the beginning is minimal due to prior length manipulation of the muscles. After 60 min of incubation, the potentiating effect of piperine was visible in both muscle types as the twitch force increased as expected (EDL: 27% [17:37]; soleus: 34% [21:47]). In the EDL muscles (Fig. 10C; Table A6), K_tr_ increased by 15% [7:22] (P < 0.001) after 60 min of incubation compared to vehicle treatment. In contrast, K_tr_ of the soleus muscles (Fig. 10D; Table A6) was unchanged after 60 min of incubation (0.5% [-0.5:1.5], P = 0.321). Hence, our data suggest that piperine increases the rate of actomyosin crossbridge cycling in EDL muscles, but not in soleus muscles.

**Figure 10.**
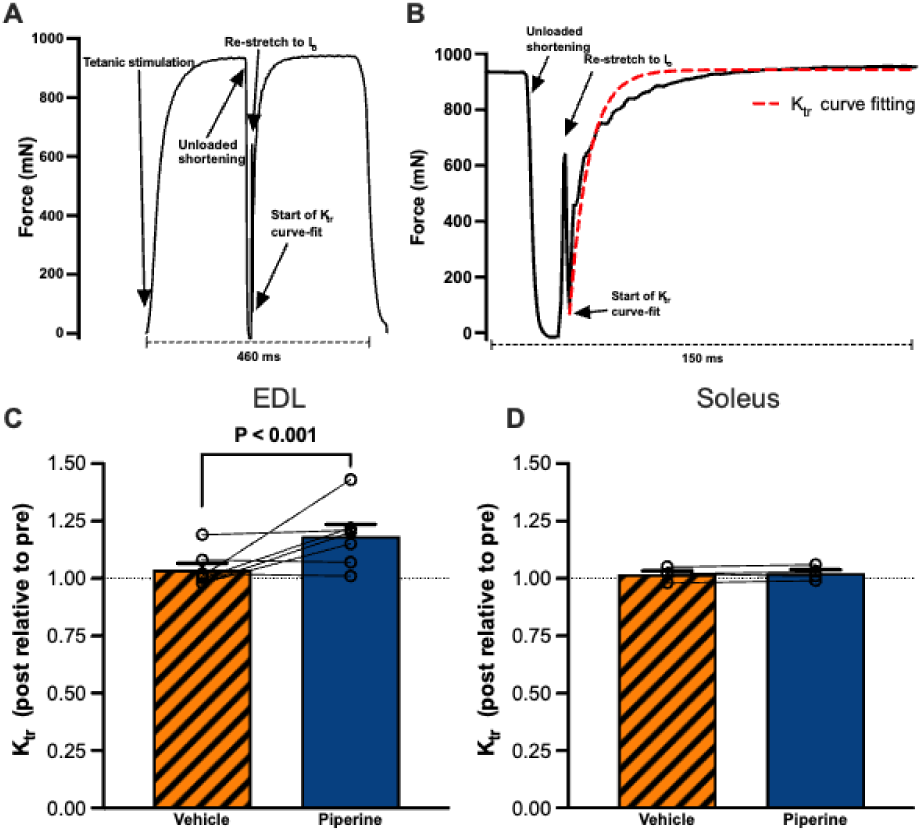
Effects of piperine on the overall actomyosin crossbridge cycling rate of intact fast (extensor digitorum longus, EDL) and slow (soleus) skeletal muscles. A, an exemplary force trace (n = 1 muscle) from an EDL muscle during a test of the isometric tension redevelopment (K_tr_). B, the insert shows the final part of the fused tetani, the unloaded shortening, the restretch, and the force redevelopment, where K_tr_ is estimated by curve-fitting (red and dotted section). C and D, the effects of 60 min of piperine (blue solid bars) and vehicle (orange shaded bars) treatment on K_tr_ normalised to the values at 0 min in EDL and soleus muscles, respectively. The P-value indicates the significance of the increase relative to baseline in piperine compared to vehicle treatment. EDL, n = 7 pairs, and soleus, n = 4 pairs. Data are shown as group means ± SEM. Circles and connecting lines represent the values of individual muscles and their contralateral pairings, respectively.

## Discussion

We show here, for the first time, that piperine induces an OFF-to-ON transition of the thick filament in resting fast and slow skeletal muscle. Further, we reveal that piperine-induced thick-filament activation leads to substantial enhancement of contractile power during low-frequency activation in both muscle types; however, only the slow muscle showed enhancement of dynamic contractility during high-frequency activation.

### Piperine induces thick-filament activation in resting skeletal muscle of both fibre types

The positioning of myosin heads relative to the thick filament governs their availability for actin interaction upon Ca²⁺-mediated thin-filament activation (Brunello & Fusi, 2024). Hence, to test if the previously observed potentiation of contractile force induced by piperine in both fibre types of skeletal muscle (Herskind *et al*., 2024) could be caused by activation of the thick filament, we applied MyoXRD on resting permeabilised muscle fibres. Taken together, our MyoXRD data show an OFF-to-ON transition of the thick filament in both fibre types in response to piperine. There was consistent directionality across all parameters, pointing to an OFF-to-ON transition; however, not all changes were statistically significant in the EDL bundles. Importantly, the regulatory protein of myosin-binding protein C (MyBP-C) has recently been shown to make a significant contribution to the M3 reflection (Koubassova *et al*., 2025). This is caused by the 3^rd^ order of the ∼43 nm MyBP-C periodicity, which overlays the ∼14.35 nm first order crown repeat, possibly hiding or exaggerating changes in myosin-head ordering on the thick filament backbone (Koubassova *et al*., 2025). Further, unpaired analyses in EDL bundles (to include non-paired data points) supported an OFF-to-ON transition by revealing significant changes (P < 0.05) in I_1.1/I1.0_, S_M3_, and S_M6,_ and borderline significance in I_M3_ (P = 0.066) and I_MLL1_ (P = 0.081) after removing an outlier from the I_M3_ (outlier > mean + 1.96 × SD). A similar approach in the soleus bundles showed significant effects (P < 0.05) on the I_1.1_/I_1.0_, S_M3_, I_M3_, S_M6_, and I_MLL1_. Since the directions of these changes were all in line with the classic OFF-to-ON signature (Linari *et al*., 2015; Ma *et al*., 2021), we find that our MyoXRD data clearly support the conclusion that piperine induces thick-filament activation in resting skeletal muscle of both fibre types.

While thick-filament activation patterns during contraction have been subject to multiple studies in recent years (Hill *et al*., 2021; Gong *et al*., 2022; Hill *et al*., 2022; Zhao *et al*., 2024; Hill *et al*., 2025), thick-filament activation in resting skeletal muscles and its consequences for contractile function have been very sparsely investigated. At present, options for studying exogenous thick-filament activation in intact skeletal muscles rely on the use of 1) transgenic animals expressing higher-than-normal levels of the thick-filament activating 2-deoxy-ATP (dATP) (Ma *et al*., 2020) or 2) cardiac-specific thick-filament activators with poor affinity for especially fast skeletal muscle (Radke *et al*., 2014; Nagy *et al*., 2015; Lindqvist *et al*., 2019; Governali *et al*., 2020; Karimi *et al*., 2024; Potoskueva *et al*., 2025). Another approach is the induction of the physiological potentiation phenomenon, post-tetanic potentiation (PTP) (Vandenboom, 2016), which potentiates force in fast-twitch fibres by Ca²^+^-dependent RLC phosphorylation. Using time-resolved MyoXRD on live tarantulas, it has been elegantly shown that PTP induces an OFF-to-ON transition of the thick filament, which causes the subsequent force potentiation (Padrón *et al*., 2020). Yet, PTP is a very short-lived force potentiation in fast skeletal muscles, which only activates the resting thick filament on a timescale of seconds. Thus, these approaches all have limitations in terms of their accessibility, ability to induce thick-filament activation acutely, or to induce activation for longer periods of time in skeletal muscle preparations. Piperine has previously been shown to activate the thick filament in resting cardiomyocytes (Jani *et al*., 2024). Our data expand on those findings, showing that piperine also induces thick-filament activation across both fibre types of skeletal muscle. Hence, piperine represents a promising tool compound for inducing thick-filament activation in skeletal muscle, offering advantages over currently limited options such as PTP, dATP, or cardiac-specific activators.

### Thick-filament activation by piperine enhances dynamic contractility in skeletal muscles, with more pronounced effects in slow muscles

Dynamic muscle function is essential for physical functioning and well-being (Freitas *et al*., 2024), making it a highly important outcome to assess when investigating modulation of the contractile machinery. We investigated the effects on dynamic muscle function in fast and slow skeletal muscles by determining force-velocity curves. Here we found that piperine-induced thick-filament activation led to a marked enhancement of V_max_ and P_max_ during low-frequency activation in both muscle types (Fig. 8A and B). These findings correlate well with the findings reported by Herskind et al. 2024 showing that piperine potentiates isometric contractions induced by low-frequency activation (Herskind *et al*., 2024). Skeletal muscles activated at low frequencies will have sub-saturating intracellular Ca^2+^ levels, and thereby the contractile machinery will be sensitive to the piperine-induced modulation of Ca^2+^-sensitivity (Herskind *et al*., 2024). In the present study, the magnitudes of potentiation of P_max_ were substantial (EDL: ∼36% and soleus: ∼54%), indicating great potential of exogenous thick-filament activation to enhance low-frequency dynamic contractility, which is the most common type of muscle contraction in vivo.

Surprisingly, we also found that piperine enhanced dynamic muscle function during high-frequency muscle activation (Fig. 9A and B). Previous studies have shown that 1) a resting pool of ON myosin motors drives low-load shortening (Linari *et al*., 2015) and 2) the number of available myosin motors governs the force-velocity curve (Piazzesi *et al*., 2007). Here we show that increasing the resting pool of ON myosin motors 1) increases V_max_ at high-frequency activation (EDL: ∼5% and soleus: ∼9%) and 2) shifts the overall force-velocity curve of both muscle types towards higher power output during high-frequency activation. In contrast to our findings, it has previously been reported that piperine does not affect the shortening velocity at 25% of maximal isometric force in maximally Ca^2+^-activated single fibres (Nogara *et al*., 2016). These experiments were performed using a temperature-jump protocol (5-25°C) while testing shortening velocity at only one external load (25% of isometric max). Hence, discrepancies between our findings and the previous study may be explained by a more comprehensive assessment of dynamic contractility performed in our study, along with differences in the experimental temperatures. Taken together, piperine seems able to modulate the underlying physiological determinants of dynamic muscle function, leading to substantial changes in dynamic contractility.

Based on our findings of the thick-filament activation induced by piperine, our contractile experiments can be viewed in a broader perspective. Hence, here we show that exogenous thick-filament activation of skeletal muscles potentiates dynamic muscle function of fast and slow muscles during both low– and high-frequency activation. Interestingly, the potential of piperine to enhance dynamic contractility during high-frequency activation seemed more pronounced in the slow muscles compared to the fast muscles, as P_max_ increased exclusively in the slow soleus muscles (∼10.5%). Thus, our data indicate fibre-type-specific differences in the level of enhancement of dynamic muscle function by exogenous thick-filament activation. These differences may be explained by recent findings, showing that the kinetics of thick-filament activation during contraction are fibre type-dependent (Gong *et al*., 2022; Hill *et al*., 2022; Zhao *et al*., 2024). Further, Hill et al. 2025 (Hill *et al*., 2025) elegantly showed that not only the kinetics, but also the overall degree of thick-filament activation during isometric fused tetani, is lower in the slow soleus compared to the fast EDL, indicating fibre-type differences in the contractile reserve capacity. In this perspective, we speculate that the large effects on dynamic muscle function in the slow soleus muscle could indicate that piperine recruits myosin heads from a pool of otherwise constitutively OFF myosin motors, representing a contractile reserve capacity which may be specifically present in slow skeletal muscle.

### Effects of piperine on force development kinetics suggest complex fibre-type-specific mechanisms

We observed seemingly contradictory effects of piperine on force development (RFD in Tables A2 and A3) and redevelopment kinetics (K_tr_ in Table A6) that reveal the complexity of piperine-potentiation across fibre types. In EDL muscles, piperine increased K_tr_ by ∼15% under conditions of maximal Ca²⁺ and a low number of attached crossbridges yet increased initial RFD of fused tetanic contractions by only ∼2%. Conversely, in soleus muscles, K_tr_ was unchanged while the initial RFD of fused tetani increased by ∼16%. In notion, Herskind et al. (Herskind *et al*., 2024) reported a similar tendency, with piperine inducing numerically larger increases in fused tetanic RFD of soleus muscles compared to EDL. The divergent findings likely reflect different rate-limiting steps in force development between fibre types and experimental conditions. K_tr_, measured after unloaded shortening and re-stretch, primarily assesses crossbridge kinetics when thin filaments are fully activated, and the number of attached crossbridges is artificially reduced. In contrast, peak RFD during normal tetanic contractions integrates multiple processes, including: 1) Ca²^+^-dependent thin– and thick-filament activation kinetics, 2) strain-dependent thick-filament activation kinetics, and 3) crossbridge cycling (Caremani *et al*., 2023; Brunello & Fusi, 2024). Also, we cannot exclude that these discrepancies may be due to isoform-dependent interactions of piperine with sarcomeric regulatory proteins, such as RLC, MyBP-C, and titin. Hence, the cause for these divergent findings may be found in the complex interaction between the effects of piperine and the effects of Ca^2+^, strain, other sarcomeric proteins, and differences between fibre types. We hypothesise that piperine has an exclusive effect on crossbridge kinetics (detected by K_tr_) in EDL muscles, which tends to support previous findings of a piperine-induced increase in resting myosin ATPase activity in fast-twitch but not in slow-twitch fibres (Nogara *et al*., 2016; Montesel *et al*., 2025). These changes were seen without substantial changes to peak RFD in EDL, which may already be near-maximal in the fast muscle type. In contrast, in the soleus muscles, piperine-induced thick-filament activation may eliminate a rate-limiting recruitment step of ON myosin during normal contractions, dramatically increasing RFD, while not affecting isolated crossbridge cycling kinetics measured by K_tr_. However, we acknowledge that these data are exploratory, and interpretation is speculative.

### Limitations

In MyoXRD experiments, we used permeabilised resting muscle fibres, a preparation known to experience cell swelling that can lead to some disruption of the resting sarcomere order (Caremani *et al*., 2021). Cell swelling can be counteracted by applying a compressor molecule such as dextran; yet, due to piperine’s poor solubility and the viscosity of dextran, we did not include dextran. However, we have no reason to believe that the initial cell swelling modifies the effect of piperine (Hessel *et al*., 2024b). Finally, we did not determine the fibre types of the fibres used in MyoXRD experiments; however, we utilised rat EDL and soleus muscles, predominantly composed of type II and I fibres, respectively (Cornachione *et al*., 2011).

Intact ex vivo muscle preparations have a limited durability of 8-9 hours in our experimental setup. In our contractile experiments, we had to apply long incubation phases to reach plateaus in force potentiation (Herskind *et al*., 2024). Further, force-velocity tests were executed throughout 60 min, making the overall experimental protocol strenuous on the preparations. We handled these issues by determining neighbouring force-velocity points at different points in the sequence to average the potential impacts of incubation time and fatigue during the force-velocity test (Kristensen *et al*., 2019). Finally, to construct force-velocity curves, we employed isotonic contractions defined as relative values of the absolute low– and high-frequency isometric maximal force at baseline. This means that changes in the contractile apparatus during the protocol manifested as changes in shortening velocity, even though the major underlying causes were likely related to changes in force output and not the ability of the sarcomere to shorten. The experimental approach and setup we applied to investigate crossbridge cycling rates K_tr_ in intact EDL and soleus muscles have previously been used by Kristensen et al. (Kristensen *et al*., 2018; Kristensen *et al*., 2019). In these studies, Kristensen et al. demonstrated that the method was sensitive to changes in K_tr_ imposed by fatigue and physiologically induced potentiation. We cannot discriminate between attachment and detachment rates by this method; therefore, we used this approach as an explorative outcome.

### Future directions

We observed marked fibre-type-dependent differences in the potential of piperine to enhance dynamic contractility. This may suggest that thick-filament activation modifies the natural contraction-induced activation kinetics of the thick filament in a fibre-type-dependent manner. Hence, future studies should utilise time-resolved MyoXRD to understand the nature of the fibre-dependent effects of exogenous thick-filament activation on dynamic muscle function. Such information will be fundamental to determining the therapeutic potential of pharmacological skeletal-muscle thick-filament activation to mitigate contractile impairments.

## Conclusion

To conclude, our data strongly indicate that piperine 1) induces thick-filament activation in resting skeletal muscle of both fibre types, and 2) enhances dynamic muscle function at submaximal activation levels in both fast and slow skeletal muscles, while only enhancing contractility of the slow muscle type during maximal activation. Hence, we propose piperine as a tool compound for the investigation of exogenous thick-filament activation in skeletal muscle. Further, we reveal that thick-filament activation in skeletal muscle leads to fibre-type-dependent magnitudes of potentiation of dynamic muscle function. Such effects must be further investigated and must be accounted for in future efforts to leverage the therapeutic potential of skeletal-muscle thick-filament activation in the treatment of contractile impairments.

## Appendix

### Effect of sarcomere length (SL) in MyoXRD experiments

To validate our MyoXRD results, we used the well-known length-dependent activation of the thick filament (Hessel *et al*., 2022) to assess the quality of our preparations. A subgroup of EDL bundles was tested at SL 2.4 and 2.7 µm before and after piperine incubation (Table A1). When comparing baseline values from the EDL bundles at SL 2.4 and SL 2.7, we saw an increase in the S_M3_ (14.36 ± 0.07 (mean ± SD) vs. 14.39 ± 0.03 nm, P = 0.027) as expected when stretching the sarcomere. From the equatorial reflections, the inter-thick-filament lattice spacing (d_10_) can be derived. When stretching EDL bundles from SL 2.4 to SL 2.7, we observed a decrease (non-significant) in d_10_ (41.41 ± 1.78 vs. 39.87 ± 2.34 nm, P = 0.121). These data suggest a decrease in d_10_ in response to stretching of the fibres, which we would expect due to compression of the sarcomere.

### Appendix Figures

**Figure A1.**
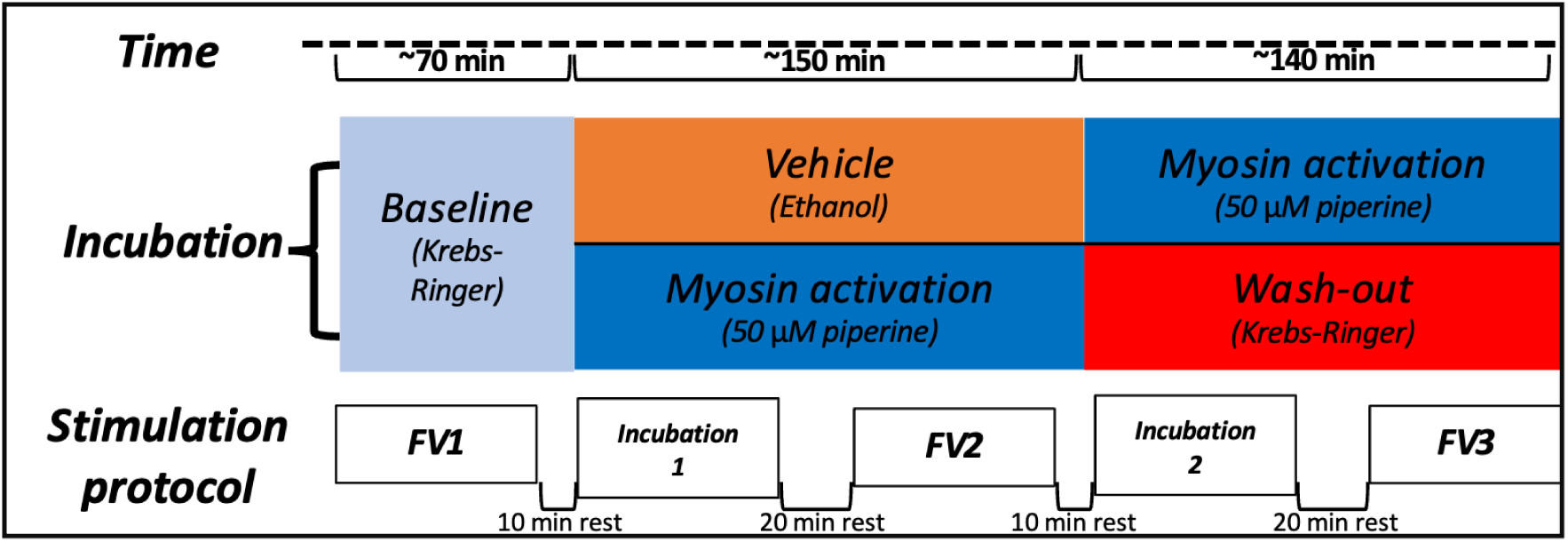
Full experimental protocol, including additional incubation and force-velocity (FV) testing. This expands on Figure 1B, which highlights the primary comparison of the first two force-velocity (FV) tests (FV1-2). Hence, an additional incubation (Incubation 2) and FV test (FV3) is shown in this figure. In the latter incubation and FV test, we investigated the ability to reverse the effects of piperine by a wash-out (substitution of KR buffer at 0, 5, 15, and 25 min) as well as the reproducibility of piperine’s effects in the contralateral muscle.

**Figure A2.**
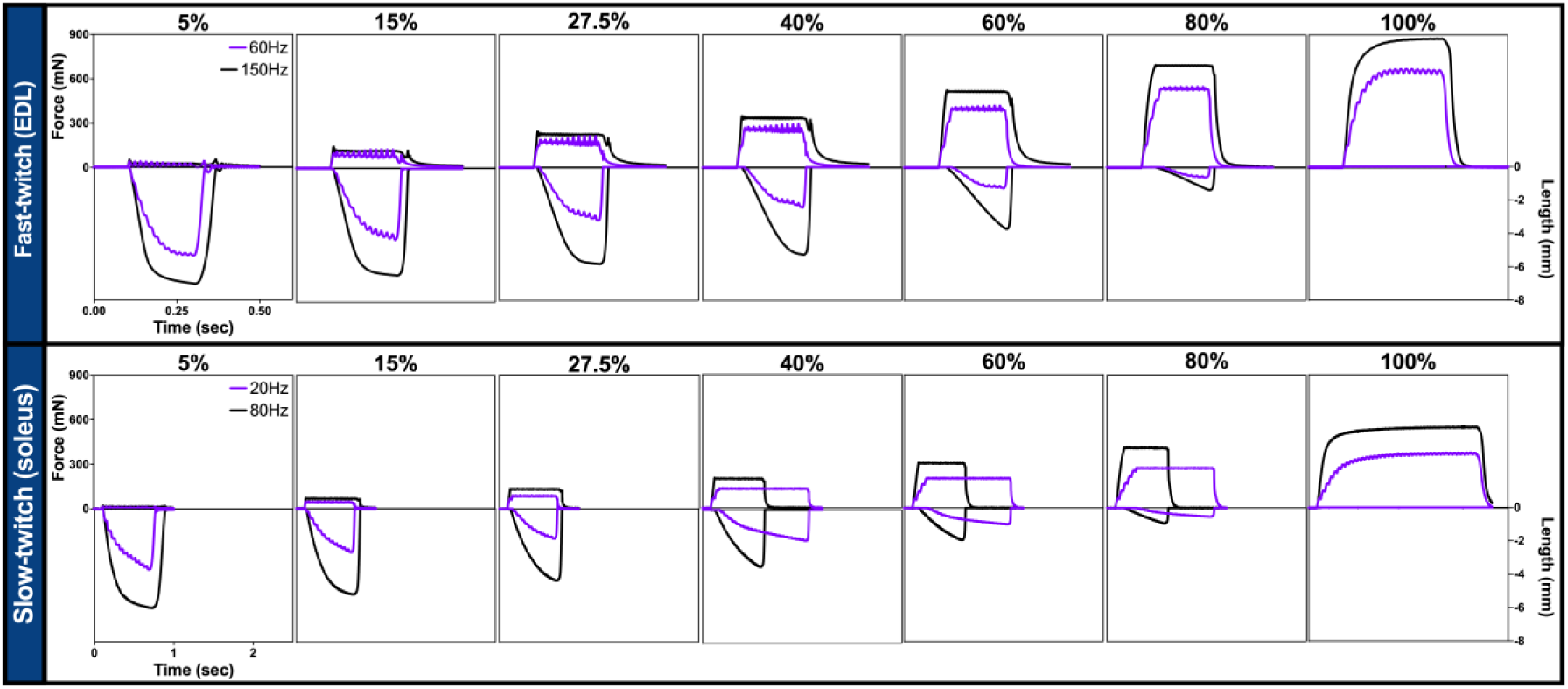
Example force– and length traces from each low– and high-frequency contraction used to construct force-velocity (FV) curves. The figure shows data from extensor digitorum longus (EDL) (n = 1) and soleus (n = 1) muscles. To determine the low– and high-frequency FV curves, each muscle performed electrically induced contractions against varying external loads (5-100% of low– and high-frequency maximal isometric force). The approach was to determine the FV curve by isotonic contractions using the afterload method. Hence, an isometric force was developed until the desired force was achieved (as a percentage of maximal isometric force), after which the muscle was allowed to shorten against the external load.

### Appendix Tables

**Table A1.** Effects of piperine on resting thick-filament structure.

|  | **EDL** |  |  |  |  |  |  | **Soleus** |  |  |
| --- | --- | --- | --- | --- | --- | --- | --- | --- | --- | --- |
|  | <u>**SL 2.4 μm**</u> |  |  | <u>**SL 2.7 μm**</u> |  |  | **Pre 2.4**<br>**vs**<br>**Pre 2.7** | <u>**SL 2.7 μm**</u> |  |  |
|  | **Pre** | **Post** | **P-value** | **Pre** | **Post** | **P-value** |  | **Pre** | **Post** | **P-value** |
| **$I_{1,1} / I_{1,0}$**<br>**(a.u.)** | 0.49 ± 0.15<br>(6) | 0.61 ± 0.25<br>(6) | 0.235<br>(5) | 0.58 ± 0.21<br>(20) | 0.94 ± 0.55<br>(14) | **0.002\***<br>(13) | 0.837<br>(3) | 0.56 ± 0.26<br>(11) | 1.31 ± 0.85<br>(14) | **<0.001\***<br>(9) |
| **$S_{M3}$**<br>**(nm)** | 14.36 ± 0.07<br>(9) | 14.37 ± 0.04<br>(7) | 0.721<br>(7) | 14.39 ± 0.03<br>(21) | 14.42 ± 0.04<br>(15) | 0.139<br>(13) | **0.027\***<br>(6) | 14.40 ± 0.03<br>(14) | 14.43 ± 0.03<br>(14) | **0.009\***<br>(11) |
| **$I_{M3}$**<br>**(a.u.)** | 522 ± 151<br>(9) | 486 ± 153<br>(7) | 0.179<br>(7) | 605 ± 166<br>(21) | 536 ± 180<br>(15) | 0.208<br>(13) | 0.294<br>(6) | 495 ± 102<br>(14) | 398 ± 141<br>(14) | **0.019\***<br>(11) |
| **$S_{M6}$**<br>**(nm)** | 7.22 ± 0.01<br>(5) | 7.22 ± 0.01<br>(3) | **0.032\***<br>(2) | 7.23 ± 0.01<br>(13) | 7.24 ± 0.01<br>(9) | **0.016\***<br>(7) | 0.121<br>(3) | 7.23 ± 0.01<br>(12) | 7.24 ± 0.01<br>(13) | 0.148<br>(8) |
| **$d_{10}$**<br>**(nm)** | 41.41 ± 1.78<br>(8) | 41.68 ± 1.64<br>(7) | 0.459<br>(7) | 39.87 ± 2.34<br>(20) | 39.17 ± 1.76<br>(13) | 0.964<br>(12) | 0.121<br>(5) | 39.75 ± 2.15<br>(11) | 40.12 ± 1.91<br>(14) | 0.070<br>(10) |
| **$I_{MLL1}$**<br>**(a.u.)** | 497 ± 296<br>(3) | 443 ± 132<br>(2) | N/a | 1663 ± 716<br>(8) | 999 ± 523<br>(5) | 0.181<br>(4) | 0.254<br>(2) | 1544 ± 665<br>(8) | 681 ± 190<br>(8) | **0.014\***<br>(6) |

**Table A2.** Effects of piperine on force and force kinetics in extensor digitorum longus (EDL) rat skeletal muscles during isometric twitch (2 Hz), unfused tetani (50 Hz), and fused tetani (150 Hz).

|  | Incubation time (min) | Stimulation frequency (Hz) | Peak force (mN) |  | RFD (mN/ms) |  | RT (ms) |  |
| --- | --- | --- | --- | --- | --- | --- | --- | --- |
|  |  |  | Vehicle (a) | Piperine (b) | Vehicle | Piperine | Vehicle | Piperine |
| Incubation 1 | 0 (pre) | 2 (twitch) | 111 ± 17 | 104 ± 18 | 6.58 ± 1.1 | 6.10 ± 1.1 | 24.3 ± 0.8 | 22.7 ± 1.0 |
|  |  | 50 (unfused) | 395 ± 72 | 363 ± 70 | 6.75 ± 1.1 | 6.40 ± 1.0 | 31.7 ± 2.1 | 30.4 ± 1.1 |
|  |  | 150 (fused) | 902 ± 153 | 908 ± 90 | 24.97 ± 4.7 | 22.54 ± 2.1 | 31.1 ± 0.6 | 29.4 ± 1.3 |
|  | 20 | 2 | 108 ± 16 | **116 ± 19\*** | 6.43 ± 1.0 | **6.70 ± 1.0\*** | **22.8 ± 0.5<sup>#</sup>** | 22.2 ± 1.2 |
|  |  | 50 | 384 ± 72 | **399 ± 76\*** | 6.62 ± 1.0 | **7.08 ± 1.0\*** | 30.5 ± 1.6 | 28.5 ± 1.2 |
|  |  | 150 | 890 ± 149* | **888 ± 90<sup>#</sup>** | 24.05 ± 4.6 | 22.66 ± 2.1 | **31.2 ± 0.8<sup>#</sup>** | 30.3 ± 1.4 |
|  | 40 | 2 | 108 ± 16 | **129 ± 17\*** | 6.42 ± 1.0 | **7.48 ± 1.0\*** | **22.6 ± 0.7<sup>#</sup>** | 21.8 ± 0.8 |
|  |  | 50 | 388 ± 72 | **454 ± 74\*** | 6.63 ± 1.0 | **7.94 ± 1.0\*** | 30.4 ± 1.7 | **28.0 ± 0.9<sup>#</sup>** |
|  |  | 150 | 879 ± 148 | 871 ± 94 | 24.07 ± 4.6 | **22.86 ± 2.1\*** | 31.3 ± 1.0 | 30.3 ± 1.5 |
|  | 60 | 2 | 108 ± 18 | **137 ± 18\*** | 6.47 ± 1.1 | **8.02 ± 1.0\*** | **22.7 ± 0.7<sup>#</sup>** | 21.9 ± 0.8 |
|  |  | 50 | 387 ± 75 | **492 ± 75\*** | 6.65 ± 1.1 | **8.48 ± 1.1\*** | 30.6 ± 1.6 | **27.5 ± 0.9<sup>#</sup>** |
|  |  | 150 | 868 ± 149 | 856 ± 101 | 24.08 ± 4.5 | **23.04 ± 2.1\*** | 31.4 ± 1.1 | 30.6 ± 1.5 |
| Assessment of dynamic muscle function including rest (90 min) |  |  |  |  |  |  |  |  |
|  |  |  | Piperine (a) | Wash-out (b) | Piperine | Wash-out | Piperine | Wash-out |
| Incubation 2 | 0 (pre) (150 min) | 2 | 101 ± 16 | **149 ± 22\*** | 6.05 ± 1.0 | **8.68 ± 1.3\*** | **22.6 ± 0.9<sup>#</sup>** | 22.6 ± 0.5 |
|  |  | 50 | 368 ± 74 | **547 ± 77\*** | 6.23 ± 1.1 | **9.18 ± 1.3\*** | 31.3 ± 1.8 | **29.4 ± 2.3<sup>#</sup>** |
|  |  | 150 | 835 ± 134 | 826 ± 93 | 23.47 ± 4.3 | **22.76 ± 2.0\*** | **31.3 ± 0.8<sup>#</sup>** | 31.6 ± 0.9 |
|  | 20 | 2 | 112 ± 18 | **136 ± 20\*** | 6.75 ± 1.1 | **7.98 ± 1.2\*** | **23.0 ± 0.3<sup>#</sup>** | 22.0 ± 0.6 |
|  |  | 50 | 404 ± 84 | **487 ± 74\*** | 6.92 ± 1.1 | **8.44 ± 1.1\*** | 30.6 ± 1.9 | 29.5 ± 1.7 |
|  |  | 150 | 813 ± 130 | 825 ± 92 | 23.47 ± 4.3 | **22.76 ± 2.1\*** | 31.8 ± 0.8 | 31.1 ± 0.7 |
|  | 40 | 2 | 124 ± 22 | **127 ± 20\*** | 7.37 ± 1.3 | **7.48 ± 1.2\*** | 23.3 ± 0.6 | 21.5 ± 0.7 |
|  |  | 50 | 460 ± 90 | **453 ± 77\*** | 7.78 ± 1.4 | **7.96 ± 1.1\*** | 29.9 ± 1.5 | 29.7 ± 1.6 |
|  |  | 150 | 808 ± 128 | 823 ± 91 | 23.58 ± 4.2 | **22.76 ± 2.1\*** | 31.8 ± 0.8 | 31.1 ± 0.8 |
|  | 60 | 2 | 134 ± 24 | 122 ± 20 | 7.90 ± 1.4 | 7.26 ± 1.2 | 23.2 ± 0.8 | 21.3 ± 0.7 |
|  |  | 50 | 495 ± 94 | 428 ± 80 | 8.45 ± 1.5 | 7.64 ± 1.1 | 29.3 ± 1.3 | 29.8 ± 1.4 |
|  |  | 150 | 804 ± 126 | 817 ± 90 | 23.70 ± 4.2 | **22.68 ± 2.1\*** | 32.3 ± 1.0 | 31.1 ± 0.7 |

The table shows force (millinewton) and force development (rate of force development (RFD)) (millinewton/millisecond) and relaxation kinetics (relaxation time (RT)) (milliseconds from 90-10% of force during relaxation) derived from EDL muscles for two 1-hour incubations. Contralateral muscles were exposed to two distinct incubation sequences (explained in Fig. A1), exposing muscles to vehicle (orange), piperine (50 µM) (blue), or a wash-out (fresh Krebs-Ringer) (red). Values in bold with \* and <sup>#</sup> indicate that muscles increased or decreased more, respectively, relatively to baseline than the opposite treatment using RM ANOVAs. n = 6 muscles in sequence a and 5 muscles in sequence b. Values are presented as mean ± standard deviation (SD).

**Table A3.** Effects of piperine on force and force kinetics in slow soleus rat skeletal muscles during isometric twitch (2 Hz), unfused tetani (20 Hz), and fused tetani (80 Hz).

|  | Incubati<br>on time<br>(min) | Stimulation<br>frequency<br>(Hz) | Peak force (mN) |  | RFD (mN/ms) |  | RT (ms) |  |
| --- | --- | --- | --- | --- | --- | --- | --- | --- |
|  |  |  | Vehicle<br>(a) | Piperine<br>(b) | Vehicle | Piperine | Vehicle | Piperine |
| Incubation 1 | 0 (pre) | 2 (twitch) | 49 ± 14 | 54 ± 14 | 1.91 ± 0.4 | 2.10 ± 0.6 | 70.6 ± 9 | 74.2 ± 3.7 |
|  |  | 20 (unfused) | 257 ± 84 | 305 ± 57 | 1.94 ± 0.4 | 2.13 ± 0.6 | 98.8 ± 13.7 | 102.7 ± 13.7 |
|  |  | 80 (fused) | 463 ± 87 | 529 ± 93 | 4.13 ± 0.9 | 4.87 ± 0.9 | 94.4 ± 14 | 97.8 ± 10.5 |
|  | 10 | 2 | 47 ± 14 | **57 ± 15\*** | 1.84 ± 0.4 | **2.20 ± 0.6\*** | 72.3 ± 8.3 | 76.1 ± 2.6 |
|  |  | 20 | 258 ± 83 | 320 ± 58 | 1.87 ± 0.4 | **2.23 ± 0.6\*** | 100.1 ± 14 | 103.8 ± 16.6 |
|  |  | 80 | 465 ± 89 | 535 ± 97 | 4.14 ± 0.9 | **5.13 ± 0.9\*** | 95.4 ± 14.2 | **94.9 ± 10.5<sup>#</sup>** |
|  | 20 | 2 | 47 ± 14 | **61 ± 16\*** | 1.84 ± 0.4 | **2.34 ± 0.7\*** | 70.4 ± 8.4 | 74.5 ± 3.1 |
|  |  | 20 | 256 ± 83 | **329 ± 57\*** | 1.88 ± 0.4 | **2.40 ± 0.6\*** | 98.7 ± 13.3 | **99 ± 16.1<sup>#</sup>** |
|  |  | 80 | 464 ± 89 | 539 ± 100 | 4.16 ± 0.9 | **5.34 ± 1.0\*** | 94.7 ± 13.6 | **92.9 ± 10<sup>#</sup>** |
|  | 30 | 2 | 47 ± 14 | **65 ± 17\*** | 1.84 ± 0.4 | **2.48 ± 0.7\*** | 70 ± 8.4 | 74.3 ± 2.9 |
|  |  | 20 | 256 ± 82 | **340 ± 57\*** | 1.85 ± 0.4 | **2.54 ± 0.7\*** | 98.1 ± 13.4 | **96 ± 15.1<sup>#</sup>** |
|  |  | 80 | 463 ± 89 | 539 ± 100 | 4.16 ± 1.0 | **5.44 ± 1.0\*** | 94.4 ± 13.4 | **92.8 ± 9.4<sup>#</sup>** |
|  | 40 | 2 | 46 ± 14 | **68 ± 17\*** | 1.82 ± 0.5 | **2.54 ± 0.7\*** | 69.8 ± 7.7 | 75.4 ± 5.3 |
|  |  | 20 | 253 ± 82 | **347 ± 57\*** | 1.85 ± 0.4 | **2.62 ± 0.7\*** | 97.8 ± 13.2 | **94.4 ± 13.8<sup>#</sup>** |
|  |  | 80 | 462 ± 90 | 537 ± 100 | 4.16 ± 1.0 | **5.52 ± 1.0\*** | 94.3 ± 13.3 | **93.6 ± 9<sup>#</sup>** |
|  | 50 | 2 | 46 ± 14 | **72 ± 20\*** | 1.81 ± 0.5 | **2.59 ± 0.8\*** | 69.8 ± 7.9 | 76.8 ± 6.7 |
|  |  | 20 | 251 ± 82 | **353 ± 57\*** | 1.82 ± 0.4 | **2.70 ± 0.7\*** | 97.9 ± 12.9 | **93.8 ± 13<sup>#</sup>** |
|  |  | 80 | 461 ± 90 | 535 ± 101 | 4.13 ± 1.0 | **5.59 ± 1.0\*** | 94.3 ± 13.2 | 95.1 ± 8.9 |
|  | 60 | 2 | 45 ± 14 | **72 ± 19\*** | 1.80 ± 0.5 | **2.64 ± 0.8\*** | **68.6 ± 7.8<sup>#</sup>** | 76.2 ± 5 |
|  |  | 20 | 248 ± 82 | **357 ± 59\*** | 1.82 ± 0.4 | **2.76 ± 0.7\*** | 97.8 ± 12.8 | **94.1 ± 12.4<sup>#</sup>** |
|  |  | 80 | 459 ± 90 | 531 ± 100 | 4.15 ± 1.0 | **5.65 ± 1.0\*** | 93.9 ± 13.1 | 96.7 ± 8.6 |
| Assessment of dynamic muscle function including rest (90 min) |  |  |  |  |  |  |  |  |
|  |  |  | Piperine<br>(a) | Wash-out<br>(b) | Piperine | Wash-out | Piperine | Wash-out |
| Incubation 2 | 0 (pre)<br>(150 min) | 2 | 44 ± 13 | **72 ± 20\*** | 1.75 ± 0.5 | **2.64 ± 0.8\*** | **69.1 ± 7.5<sup>#</sup>** | 79.5 ± 6.6 |
|  |  | 20 | 242 ± 80 | **366 ± 62\*** | 1.81 ± 0.4 | **2.79 ± 0.8\*** | 98.4 ± 14.1 | 101.5 ± 16 |
|  |  | 80 | 438 ± 100 | 510 ± 99 | 4.08 ± 1.1 | **5.44 ± 1.0\*** | **95.2 ± 15.2<sup>#</sup>** | 106.8 ± 10.3 |
|  | 10 | 2 | 49 ± 14 | **67 ± 20\*** | 1.90 ± 0.5 | **2.46 ± 0.8\*** | **70.2 ± 8<sup>#</sup>** | 81.6 ± 7.5 |
|  |  | 20 | 250 ± 77 | **354 ± 63\*** | 1.92 ± 0.5 | **2.57 ± 0.7\*** | **94.5 ± 13.3<sup>#</sup>** | 101.7 ± 15.3 |
|  |  | 80 | 437 ± 101 | 510 ± 99 | 4.31 ± 1.1 | **5.30 ± 1.1\*** | **89.2 ± 13.1<sup>#</sup>** | 107.2 ± 10.5 |
|  | 20 | 2 | 54 ± 15 | 63 ± 19 | 2.09 ± 0.5 | 2.36 ± 0.8 | 70.8 ± 7 | 77.7 ± 7 |
|  |  | 20 | 262 ± 77 | **338 ± 61\*** | 2.11 ± 0.5 | **2.45 ± 0.8\*** | **91.8 ± 12.8<sup>#</sup>** | 100.5 ± 14.5 |
|  |  | 80 | 457 ± 119 | 508 ± 98 | 4.46 ± 1.2 | 5.16 ± 1.0 | **88.8 ± 11.7<sup>#</sup>** | 104.9 ± 10.1 |
|  | 30 | 2 | 58 ± 15 | 59 ± 19 | **2.19 ± 0.6\*** | 2.22 ± 0.8 | 71.5 ± 7.3 | 77.1 ± 5.9 |
|  |  | 20 | 273 ± 77 | 331 ± 58 | **2.21 ± 0.5\*** | 2.32 ± 0.8 | **90.1 ± 11.8<sup>#</sup>** | 99.4 ± 13.4 |
|  |  | 80 | 440 ± 101 | 507 ± 98 | **4.56 ± 1.2\*** | 5.05 ± 1.0 | **89.8 ± 11.2<sup>#</sup>** | 103.8 ± 9.4 |
|  | 40 | 2 | **60 ± 16\*** | 58 ± 19 | **2.24 ± 0.5\*** | 2.19 ± 0.8 | 70.7 ± 8.1 | 74.3 ± 6.7 |
|  |  | 20 | 279 ± 77 | 320 ± 59 | **2.30 ± 0.5\*** | 2.26 ± 0.7 | **89.8 ± 11.4<sup>#</sup>** | 99.2 ± 14.1 |
|  |  | 80 | 437 ± 99 | 507 ± 97 | **4.64 ± 1.3\*** | 4.98 ± 1.0 | **91.3 ± 10.8<sup>#</sup>** | 102.4 ± 8.7 |
|  | 50 | 2 | **61 ± 18\*** | 56 ± 18 | **2.26 ± 0.6\*** | 2.18 ± 0.7 | 72.7 ± 7.6 | **72.3 ± 7<sup>#</sup>** |
|  |  | 20 | **284 ± 78\*** | 314 ± 58 | **2.33 ± 0.6\*** | 2.23 ± 0.7 | **90.3 ± 11.2<sup>#</sup>** | 99.1 ± 14.1 |
|  |  | 80 | 434 ± 99 | 507 ± 95 | **4.67 ± 1.3\*** | 4.92 ± 1.0 | 94.3 ± 12.5 | 101.2 ± 8.3 |
|  | 60 | 2 | **62 ± 18\*** | 54 ± 18 | **2.29 ± 0.6\*** | 2.12 ± 0.7 | 73.4 ± 7.8 | **72.2 ± 5.7<sup>#</sup>** |
|  |  | 20 | **286 ± 79\*** | 306 ± 57 | **2.36 ± 0.6\*** | 2.16 ± 0.7 | **91.3 ± 11.1<sup>#</sup>** | 97.9 ± 13.5 |
|  |  | 80 | 430 ± 99 | 506 ± 94 | **4.66 ± 1.3\*** | 4.84 ± 1.0 | **94.4 ± 10.2<sup>#</sup>** | 100.4 ± 8 |

Similar to Table A2 above, the table shows force and force development and relaxation kinetics from two 1-hour incubations (Fig. A1), however, for the soleus muscles. Values in bold with \* and # indicate that muscles increased or decreased more, respectively, relatively to baseline than the opposite treatment ( $P \leq 0.05$ ) using RM ANOVAs. $n = 6$ muscles in sequences a and b, respectively. Values are presented as mean ± SD.

**Table A4.**
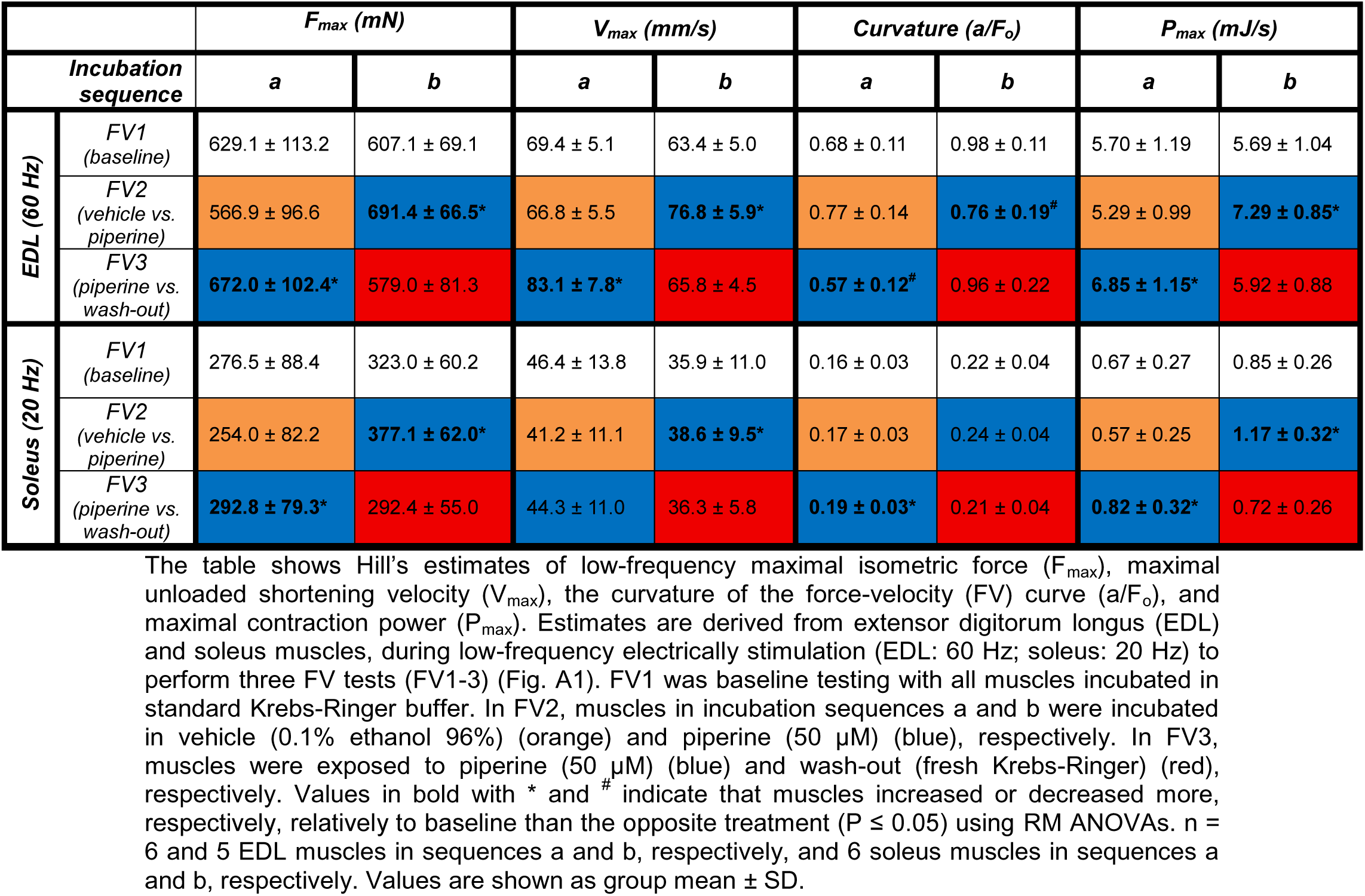
Absolute Hill estimates from the three low-frequency force-velocity tests (FV1-3).

| | | $F_{max}$ (mN) | | $V_{max}$ (mm/s) | | Curvature ($a/F_o$) | | $P_{max}$ (mJ/s) | |
| --- | --- | --- | --- | --- | --- | --- | --- | --- | --- |
| Incubation sequence |  | *a* | *b* | *a* | *b* | *a* | *b* | *a* | *b* |
| EDL (60 Hz) | FV1 (baseline) | 629.1 ± 113.2 | 607.1 ± 69.1 | 69.4 ± 5.1 | 63.4 ± 5.0 | 0.68 ± 0.11 | 0.98 ± 0.11 | 5.70 ± 1.19 | 5.69 ± 1.04 |
|  | FV2 (vehicle vs. piperine) | 566.9 ± 96.6 | **691.4 ± 66.5\*** | 66.8 ± 5.5 | **76.8 ± 5.9\*** | 0.77 ± 0.14 | **0.76 ± 0.19<sup>#</sup>** | 5.29 ± 0.99 | **7.29 ± 0.85\*** |
|  | FV3 (piperine vs. wash-out) | **672.0 ± 102.4\*** | 579.0 ± 81.3 | **83.1 ± 7.8\*** | 65.8 ± 4.5 | **0.57 ± 0.12<sup>#</sup>** | 0.96 ± 0.22 | **6.85 ± 1.15\*** | 5.92 ± 0.88 |
| Soleus (20 Hz) | FV1 (baseline) | 276.5 ± 88.4 | 323.0 ± 60.2 | 46.4 ± 13.8 | 35.9 ± 11.0 | 0.16 ± 0.03 | 0.22 ± 0.04 | 0.67 ± 0.27 | 0.85 ± 0.26 |
|  | FV2 (vehicle vs. piperine) | 254.0 ± 82.2 | **377.1 ± 62.0\*** | 41.2 ± 11.1 | **38.6 ± 9.5\*** | 0.17 ± 0.03 | 0.24 ± 0.04 | 0.57 ± 0.25 | **1.17 ± 0.32\*** |
|  | FV3 (piperine vs. wash-out) | **292.8 ± 79.3\*** | 292.4 ± 55.0 | 44.3 ± 11.0 | 36.3 ± 5.8 | **0.19 ± 0.03\*** | 0.21 ± 0.04 | **0.82 ± 0.32\*** | 0.72 ± 0.26 |

The table shows Hill's estimates of low-frequency maximal isometric force ( $F_{max}$ ), maximal unloaded shortening velocity ( $V_{max}$ ), the curvature of the force-velocity (FV) curve ( $a/F_o$ ), and maximal contraction power ( $P_{max}$ ). Estimates are derived from extensor digitorum longus (EDL) and soleus muscles, during low-frequency electrically stimulation (EDL: 60 Hz; soleus: 20 Hz) to perform three FV tests (FV1-3) (Fig. A1). FV1 was baseline testing with all muscles incubated in standard Krebs-Ringer buffer. In FV2, muscles in incubation sequences a and b were incubated in vehicle (0.1% ethanol 96%) (orange) and piperine (50 μM) (blue), respectively. In FV3, muscles were exposed to piperine (50 μM) (blue) and wash-out (fresh Krebs-Ringer) (red), respectively. Values in bold with \* and <sup>#</sup> indicate that muscles increased or decreased more, respectively, relatively to baseline than the opposite treatment ( $P \leq 0.05$ ) using RM ANOVAs. $n =$ 6 and 5 EDL muscles in sequences a and b, respectively, and 6 soleus muscles in sequences a and b, respectively. Values are shown as group mean ± SD.

**Table A5.** Absolute Hill estimates from the three high-frequency force-velocity tests (FV1-3).

| | | $F_{max}$ (mN) | | $V_{max}$ (mm/s) | | Curvature ($a/F_0$) | | $P_{max}$ (mJ/s) | |
| --- | --- | --- | --- | --- | --- | --- | --- | --- | --- |
| Incubation sequence |  | a | b | a | b | a | B | a | b |
| EDL (150 Hz) | FV1 (baseline) | 937.1 ± 115.9 | 955.4 ± 88.6 | 135.8 ± 3.7 | 132.1 ± 4.2 | 0.39 ± 0.04 | 0.45 ± 0.03 | 14.55 ± 2.93 | 14.94 ± 1.98 |
|  | FV2 (vehicle vs. piperine) | 851.7 ± 138.6 | 848.4 ± 99.8 | 134.2 ± 3.3 | **137.9 ± 2.8\*** | 0.44 ± 0.03 | **0.44 ± 0.02<sup>#</sup>** | 13.88 ± 2.60 | 13.96 ± 1.79 |
|  | FV3 (piperine vs. wash-out) | 797.8 ± 119.8 | 818.4 ± 90.4 | 138.8 ± 4.0 | 133.7 ± 3.2 | 0.42 ± 0.06 | 0.48 ± 0.05 | 13.16 ± 2.30 | 13.64 ± 1.64 |
| Soleus (80 Hz) | FV1 (baseline) | 464.7 ± 98.6 | 563.0 ± 93.6 | 51.5 ± 6.3 | 53.6 ± 12.6 | 0.16 ± 0.02 | 0.20 ± 0.02 | 1.82 ± 0.59 | 2.40 ± 0.61 |
|  | FV2 (vehicle vs. piperine) | 443.1 ± 97.0 | 515.0 ± 98.5 | 51.3 ± 7.7 | **58.0 ± 12.8\*** | 0.17 ± 0.02 | 0.19 ± 0.03 | 1.75 ± 0.59 | **2.54 ± 0.53\*** |
|  | FV3 (piperine vs. wash-out) | 408.2 ± 92.6 | 501.9 ± 85.5 | 55.5 ± 8.4 | 54.7 ± 11.6 | **0.17 ± 0.01\*** | 0.19 ± 0.03 | 1.83 ± 0.64 | 2.26 ± 0.60 |

Similar to Table A4 above, the table shows Hill's estimates of $F_{max}$ , $V_{max}$ , the curvature of the FV curve, and $P_{max}$ . In this table, estimates are derived from extensor digitorum longus (EDL) and soleus muscles, during high-frequency stimulation (EDL: 150 Hz; soleus: 80 Hz). The setup and colour-coding are identical to Table A4 above. Values in bold with \* and # indicate that muscles increased or decreased more, respectively, relatively to baseline than the opposite treatment ( $P \leq$ 0.05) using paired two-way ANOVAs. $n = 6$ and 5 EDL muscles in sequences a and b, respectively, and 6 soleus muscles in sequences a and b, respectively. Values are shown as group mean ± SD.

**Table A6.** Effects of piperine and vehicle treatment on crossbridge cycling rates in intact rat extensor digitorum longus (EDL) and soleus muscles.

|  | *Incubation time*<br>*(min)* | *K<sub>tr</sub> (s<sup>-1</sup>)* |  |
| --- | --- | --- | --- |
|  |  | *Vehicle* | *Piperine* |
| **EDL** | 0 (pre) | 98.4 ± 36.8 | 99.7 ± 25.5 |
|  | 15 | 99.2 ± 37.3 | 102.8 ± 27.6 |
|  | 30 | 100.1 ± 40.3 | **110.8 ± 30.3\*** |
|  | 45 | 100.3 ± 39.2 | **112.9 ± 23.3\*** |
|  | 60 | 101.9 ± 40.7 | **114.1 ± 33.0\*** |
| **Soleus** | 0 (pre) | 19.7 ± 2.5 | 18.9 ± 1.4 |
|  | 15 | 19.7 ± 2.3 | 19.1 ± 1.1 |
|  | 30 | 19.8 ± 2.2 | 19.2 ± 1.0 |
|  | 45 | 19.8 ± 2.3 | 19.2 ± 1.0 |
|  | 60 | 19.9 ± 2.2 | 19.3 ± 1.0 |

The table shows the absolute values of the rate constant of force redevelopment ( $K_{tr}$ ). Contralateral muscles were exposed to piperine (50 μM) or vehicle (ethanol) treatment for 60 min, with outcomes being assessed every 15<sup>th</sup> min. Values in bold with \* indicate that muscles increased more relative to baseline than the opposite treatment at the given timepoint ( $P \leq 0.05$ ) using 2-way ANOVAs with Tukey's correction. EDL: Time: $P = 0.002$ , Incubation: $P = 0.088$ , Time × Incubation: $P = 0.022$ . Soleus: Time: $P = 0.225$ , Incubation: $P = 0.384$ , Time × Incubation: $P = 0.577$ . $N = 7$ pairs of EDL muscles and 4 pairs of soleus muscles. Values are presented as mean ± standard deviation (SD).

## Additional Information

### Data availability statement

The data that support the findings of this study are available in the supporting information and upon request from the corresponding author.

### Author Contributions

D.Z.K., K.O., and A.L.H. conceived the experiments, and D.Z.K., K.O., J.H., M.N.K., A.J.K., and A.L.H. carried them out. D.Z.K. and A.L.H. analysed data. All authors contributed to the interpretation of the results. D.Z.K. wrote the first draft, and all authors helped shape the paper.

## Acknowledgments

We thank Khoi Nguyen from Accelerated Muscle Biotechnologies (AMB) for his assistance in the MyoXRD experiments. Also, we thank Cuno Rasmussen for his assistance in designing software for the analysis of the contractile data. We acknowledge DESY (Hamburg, Germany), a member of the Helmholtz Association HGF, for the provision of experimental facilities. Parts of this research were carried out at Petra III. Data was collected using the P03 beamline 18ID operated/provided by DESY Photon Science. We would like to thank Matthias Schwartzkopf and Jan Rubeck for assistance during the experiments. Beamtime was allocated for proposals I-20221421 and II-20241178.

